# RPGR regulates motile cilia by interfering with actin dynamics

**DOI:** 10.1101/2025.03.13.643176

**Authors:** Yang Wu, Erika Tavares, Binrun Liang, Wallace Wee, Vito Mennella, Hanchao Feng, Jiaying Cao, Jiayi Zheng, Mu He, Kirk Stephenson, Liran Hanan, Janice Min Li, Sharon D Dell, Elise Heon, Zhen Liu

## Abstract

Cilia are highly conserved cellular organelles extruding from the surface of cell types carrying either sensory (signaling) or motile functions. These include photoreceptor cells and airway epithelia, where they function in light sensation and mucociliary clearance respectively. Retinitis pigmentosa GTPase regulator (*RPGR*) variants affect both photoreceptor sensory cilia and airway motile cilia, leading to retinitis pigmentosa (RP) and in some cases, primary ciliary dyskinesia (PCD), both debilitating conditions. Not all patients develop PCD and it is unclear which *RPGR* variants predispose patients to PCD and why this happens. In this study, using nasal biopsy samples of patients with *RPGR*-related RP, we leverage 2D organoid cell culturing, super-resolution microscopy, and live cell imaging to characterize the multiciliated cells from patients with different *RPGR* variants, healthy human nasal and bronchial multiciliated cells with CRISPR-modified *RPGR* function. We demonstrate for the first time that multiciliated cells with *RPGR* variants may have reduced ciliation, shorter cilia, significantly impaired cilia beat, or cilia beat incoordination, which could lead to defective mucociliary clearance and lung disease. In addition, we show the regulation of motile cilia by RPGR involves F-actin, as evidenced by temporarily reduced Gelsolin and undissolved condensed actin meshwork at the apical surface of RPGR-deficient multiciliated cells. In support, we show that the motile cilia defect can be ameliorated by treating with the actin polymerization inhibitor Latrunculin A. Though PCD was observed only in patients with variants that affect both main isoforms (*RPGR^1-19^* and *RPGR*^ORF15^), patients with variants affecting only *RPGR^ORF15^* also showed cilia and airway anomalies. Though all *RPGR* variants affected motile cilia in one way or another, *RPGR* loss of function variants affecting both isoforms are associated with more severe cilia and systemic phenotypes, the mechanisms of which involve the accumulation of apical F-actin.

**One Sentence Summary:** Loss of RPGR, through the mechanism of apical F-actin accumulation in airway multiciliated cells, leads to reduced ciliation with short cilia that have an impaired beat, leading to defective mucociliary clearance.

## INTRODUCTION

Cilia are highly conserved organelles protruding from the cell surface and are essential for cellular sensing, signaling, or motility (*1, 2*). Cilia can be generally classified into motile cilia or sensory (primary) cilia (*3*), based on their structure and whether they beat or not (Fig.1A). Motile cilia align along the surface of the respiratory tract, ependyma, and fallopian tube, beat in synchrony and function in airway mucociliary clearance, cerebral fluid circulation, and egg delivery (*1*). The sensory cilia are solitary and appear on nearly all cells, receiving environmental stimuli and eliciting signaling pathways such as the Hedgehog signaling (*4*). One example of primary cilia is the photoreceptor outer segment, a specialized sensory cilium dedicated to phototransduction (*5*).

**Fig. 1.**
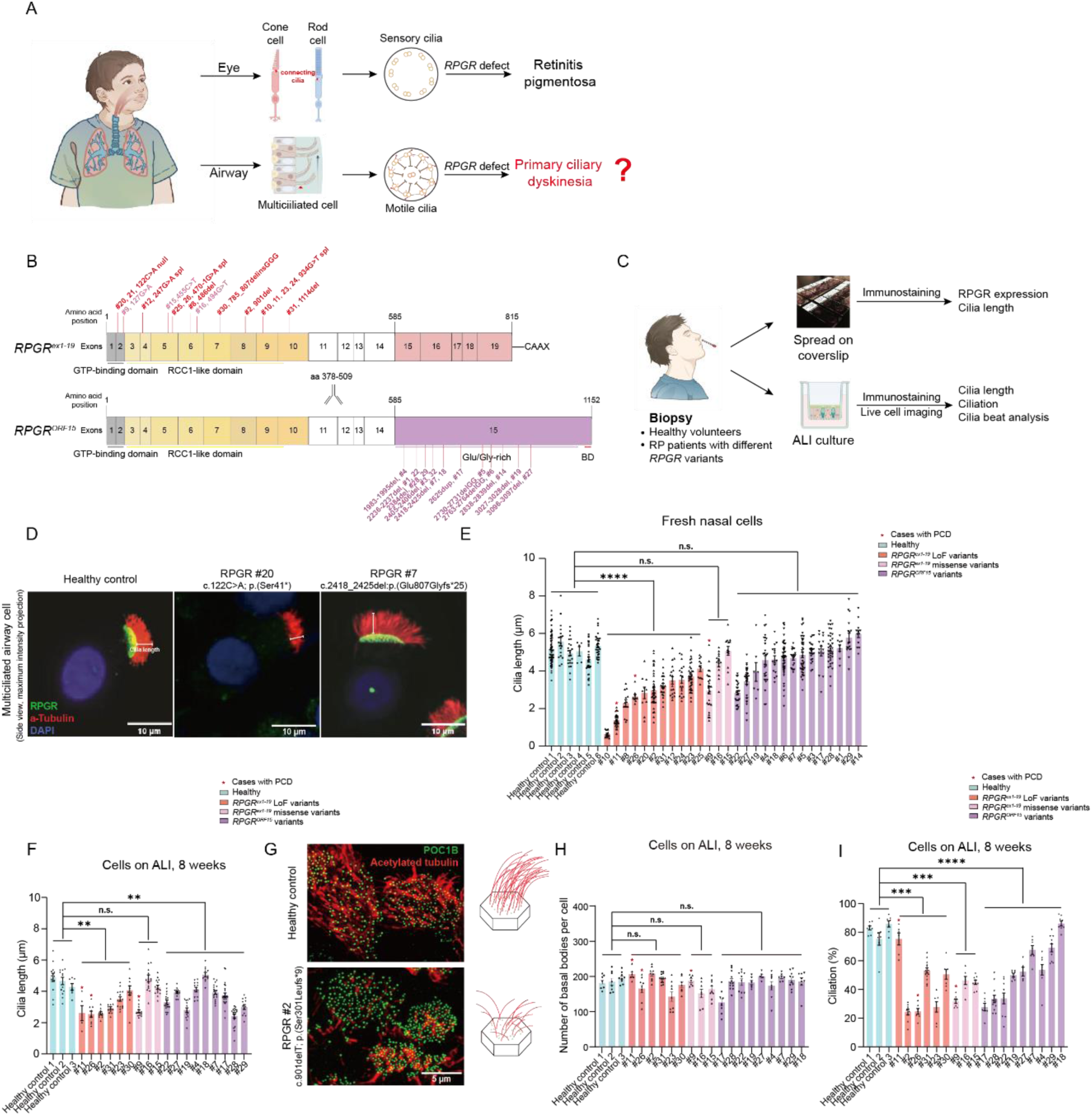
Multiciliated nasal cells (MCCs) with pathological *RPGR* variants present with short cilia, and decreased ciliation level. **(A)** Cartoon showing *RPGR* as the causative gene for RP and PCD as well as the focus of this study. **(B)** Distributions of *RPGR* variants of the 32 patients with RP participating in this study. Red indicates variants that affect both isoforms and correspond to loss of function or splicing; pink correspond to variants with missense effect; and purple correspond to loss of function variants affecting only *RPGR^ORF15^*. Variants of case # 9 and 15 are predicted to be missense and affect splicing. **(C)** The proposed workflow for studying nasal cilia structure and function related to *RPGR* variants. **(D)** Immunostaining of human nasal cells of controls and patients with *RPGR* variants. RPGR antibodies (green) and acetylated tubulin antibodies (red) staining showed reduced cilia length in certain MCCs of RP patients. **(E)** Cilia length measurements for the nasal MCCs from both controls and RP patients spread on the coverslips. the red star means the individuals with PCD. n>7 cells per sample. **(F)** Cilia length measurements for the MCCs from both controls and RP patients cultured at air-liquid interface for 8 weeks. n>6 cells per sample. **(G)** Immunostaining of basal body marker Poc1b and cilia marker acetylated tubulin showed reduced ciliation in the MCCs of a RP patient with *RPGR* damaging variant (#2, c.901delT;p.(Ser301Leufs*9)). **(H)** The number of basal bodies was unchanged for most MCCs bearing damaging variants. **(I)** Ciliation was reduced for the MCCs bearing pathological RPGR variants in either the ORF region or non-ORF15 region. n>6 cells per sample. All data are presented as average ± SEM. n.s., no significance, *, p < 0.05, **, p < 0.01, ***, p < 0.001, ****, p < 0.0001 by two-tailed t-test (E, F, H, I).

Conditions due to defective function, development, or maintenance of cilia are collectively referred to as ciliopathies. Primary ciliary dyskinesia (PCD) is an important motile ciliopathy (*6, 7*), affecting ∼1:7,500 people worldwide (*8*). Major features of PCD include oto-sino-pulmonary diseases which can be life-threatening because of deterioration of lung capacity. Retinitis pigmentosa (RP) is a genetically heterogeneous disorder characterized by degeneration of photoreceptors, leading to night blindness, tunnel vision, and finally sight loss (*9*). The incidence rate of RP ranges between 1 in 2,500-7,000 (*10*). Variants in genes associated with primary cilia can also cause RP (*11*).

Deleterious variants involving the *Retinitis pigmentosa GTPase regulator* (*RPGR*) gene account for ∼70% of X-linked RP (*12–14*) and are also a known cause of X-linked PCD (*15*). *RPGR* is expressed in both sensory and motile cilia (*16, 17*). As a result, patients with *RPGR* deleterious variants could present both motile and sensory ciliopathy phenotypes (Fig. 1A). *RPGR* is expressed in two major isoforms (Fig. 1B) (*18–21*), one is constitutively expressed in multiple organs and cell types (NM_000328.3; *RPGR^ex1-19^*) and the other has a restricted expression pattern that includes the retina (NM_001034853.2; *RPGR^ORF15^*). *RPGR^ex1-19^* has 19 exons of which *RPGR^ORF15^* shares exon 1-14 (*18, 22–25*). *RPGR^ORF15^*is an X-linked RP mutational hotspot and is predominantly expressed in the retina, while *RPGR^ex1-19^* is ubiquitously expressed and was initially regarded as the sole isoform in airway multiciliated cells (MCCs) (*17, 18*). In the retina, RPGR contributes to outer segment membrane disk renewal and variants involving both *RPGR^ex1-19^* and *RPGR^ORF15^* contribute to early onset RP (*26*). However, RPGR’s role in motile cilia remains unclear. While most PCD causative genes encode either axoneme components, dynein arm docking/assembly factors, or transcription factors governing the multiciliogenesis program, *RPGR* is different and maybe involved in ciliary transport (*27–30*). The gold standard for confirming PCD diagnosis is either ciliary structural defects detected by TEM and/or biallelic mutations in the known PCD genes detected by DNA sequencing. However, this standard cannot diagnose all PCD cases including *RPGR* variants as *RPGR* variants could be of unknown significance and don’t usually lead to TEM changes (*31, 32*). There have been clinical reports showing that variants in *RPGR* cause PCD (*15, 19, 33, 34*) (Table S1, https://www.ncbi.nlm.nih.gov/clinvar) and that these patients showed static cilia, impaired cilia beat, or disrupted beat coordination (*33*). However, these were mostly case studies based on a limited number of patients. Which variants predispose patients to PCD and how RPGR regulates motile cilia properties is still unclear. This knowledge gap can delay the clinical diagnosis of PCD until damage occurs and pulmonary function is affected, which can affect patient outcomes.

Using super-resolution microscopy and live cell cilia analysis of patient samples, we observed that patients with *RPGR* pathological variants (variants before exon 14 will be referred to as *RPGR* variants) presented sparse, short, and mostly static motile cilia in human nasal epithelial cells (HNCs). These phenotypes were recapitulated in CRISPR-mediated *RPGR* KO MCCs. For *RPGR^ORF15^* variants, while nasal cilia length was not different from healthy controls, they were sparser and grew slower for MCCs cultured at the air-liquid interface. For the cilia beating of MCCs bearing *RPGR^ORF15^* variants, most cilia exhibited decreased motility, but the beat amplitude and waveform were all abnormal and the beat coordination within and between cells was disrupted.

Investigating the mechanisms underlying how RPGR regulates motile cilia revealed its role in regulating F-actin dynamics. At the apical surface of MCCs with *RPGR* loss of function variants, a condensed actin meshwork accumulates, preventing some cilia from extruding from the surface, and restricting cilia movement on the cell surface. The actin meshwork is possibly caused by a temporarily reduced level of Gelsolin at the apical surface. To examine whether the phenotype was specific to F-actin changes, we treated MCCs bearing *RPGR* variants with Latrunculin A, an actin polymerization inhibitor, and found that, similar to sensory cilia in RPE-1 cells, cilia length and ciliation defect could be ameliorated, and cilia beat could be partially restored.

To summarize, using a cohort of 32 patients (Table S1) with variants in *RPGR* and *RPGR^ORF15^*, we provide the first mechanistic study of RPGR’s function in the respiratory system. We found that although symptoms of PCD are more prevalent with *RPGR* variants, cilia anomalies are seen in both isoforms, and patients with *RPGR^ORF15^* variants have some respiratory findings, though not characteristic of PCD (Table S2). The more severe cilia phenotypes were associated with *RPGR* loss of function variants. This work also reveals a different role of *RPGR* compared to other PCD causative genes: RPGR tunes F-actin dynamics at the apical surface to control multiciliogenesis and to regulate cilia beat machinery. The methods reported here can be translated clinically to identify individuals at risk of mucociliary clearance defects and improve patient outcomes with appropriate management.

## RESULTS

### Patient characterization

We recruited a cohort of 32 RP patients with a wide range of RPGR variants including one female with retinal degeneration (case 21) (Fig. 1C, Table S1), and six healthy volunteers. All patients were recruited through eye clinics and had retinal degeneration. Fresh nasal cilia were available in 29 cases for cilia analysis. The mean age at the time of nasal cilia biopsy was 30 years (range 8-63 years). The pathogenicity of the variants identified was validated using available pathogenicity and conservation predictive algorithms, allele frequency databases, and splicing assays (*35–40*) (Table S1, Fig.S1). Variants affecting both isoforms were found in 16 patients: 8 were considered loss of function (RPGR LOF), 5 were missense resulting in an in-frame deletion (RPGR in frame deletion), two had missense variants with loss of transcript due to incomplete splicing (RPGR missense splicing), and one had a missense (RPGR missense). Loss of function variants affecting the RPGRORF15 isoform accounted for the remaining 16 patients (Table S1).

Respiratory manifestations of PCD include neonatal respiratory distress, chronic nasal congestion, chronic cough and wheezing, sinusitis, bronchitis, pneumonia, and/or otitis media. Clinical assessments of PCD patients show obstructive lung disease by lung function test, bronchiectasis by chest CT, chronic abnormalities by chest radiograph, and/or a low exhaled nitric oxide level (< 77 nL/min). Of the 32 patients in this study, 6 were highly suspicious of PCD (severe respiratory changes with reduced nNO and/or chest X-ray/CT anomalies), all of which had RPGR variants. Of the 32 patients recruited, 19 out of 31 had sinus diseases, 15 out of 30 had lower airway diseases, and 19 patients presented ear issues including hearing loss, and/or otitis media (Table S2). For the clinical tests, 9 out of 28 patients had obstructive lung functions and 11 out of 27 patients had Chest X-ray and/or CT anomalies. Of the 16 cases carrying ORF15 variants, 5 had sinus diseases, 4 had lower airway diseases, 8 had hearing issues including 2 with hearing loss. Clinical tests suggested 2 had obstructive airway diseases and 3 had abnormal chest radiographs.

### Patients with pathological *RPGR* variants show cilia defects

Fresh MCCs isolated from 6 healthy volunteers and 29 RP patients were immunostained with RPGR antibodies and antibodies targeting the cilia marker acetylated tubulin for either 3D-SIM or Confocal microscopy imaging. RPGR was absent from most patients with variants that affect both isoforms but was properly localized in the cells bearing *RPGR^ORF15^* variants (Fig. 1D). Cilia length measurements showed that for most patients with *RPGR* loss of function and in frame deletion variants, cilia are significantly shorter (p < 0.0001, N=11) while patients with *RPGR^ORF15^* variants (n.s., N=14) displayed normal cilia length (Fig. 1E). This suggests that overall RPGR is important for motile cilia growth or length regulation.

Interestingly, in a female patient (Case #21) with severe RP due to *RPGR* variant (c.122C>A; p. (Ser41*), around half of her MCCs did not express RPGR and presented short cilia because of unfavorable X-inactivation (Fig. S2A). Thus, within the same biological background, the cells with shorter cilia were the ones lacking RPGR.

Because fresh nasal cells might be subject to infections and other environmental stressors, we re-differentiated the isolated basal cells on the air-liquid interface (ALI) for further assessment. Such differentiated cells were available for 18 cases. We measured the cilia length for the differentiated cells at 4 weeks (Fig. S2B) and 8 weeks (Fig. 1F) using the signal of acetylated tubulin. Six of nine patients (66%) with variants in *RPGR* and 8 out of 9 patients with *RPGR^ORF15^* variants have significantly shorter motile cilia than control individuals (p<0.0001 and 0.05 respectively). The shorter cilia length from MCCs bearing *RPGR^ORF15^* variants suggests that *RPGR^ORF15^* may be important for the growth of cilia. The ALI cell culture also allowed us to quantify the ciliation level defined by the number of cilia divided by the number of basal bodies per MCC (Fig. 1G-I). For patients with either *RPGR* (p<0.0001, N=9) or *RPGR^ORF15^* (8/9, p<0.05, N=9) variants, the ciliation level was significantly lower than healthy controls (Fig. 1I).

Altogether, our characterization of the MCCs of patients with a wide variety of pathogenic and likely pathogenic *RPGR* variants shows that disruption of RPGR function may lead to reduced ciliation (8/9 for *RPGR* variants and 8/9 for *RPGR^ORF15^*variants) and shortened cilia length(13/14 for *RPGR* variants and 7/14 for *RPGR^ORF15^* variants, nasal cilia).

### Damaging *RPGR* variants affect cilia beat

To investigate how RPGR regulates cilia motility, cilia beating was measured on MCCs re-differentiated at ALI (8 weeks). We stained the whole filter with wheat germ agglutinin (WGA)-Alexa 488 to mark the highly glycosylated motile cilia (*41, 42*) and used a method developed for automatic high throughput measurement for each entire field of view to avoid observation bias (Fig. 2A). The frequency of pixel signal fluctuation of the collected cilia beat video was used to first represent the overall motility information in the field of interest (Fig. 2B and S3). For healthy control cells, motile cilia beat had a distinguishable power stroke, recovery stroke as well as synchronization of metachronal wave (Video S1). On the contrary, MCCs from RP patients showed noticeable heterogeneity in those features, between patients as well as among cells of the same patient (Video S1). To quantify the differences, we first measured the cilia beat frequency for individual cells from patients with different *RPGR* variants. Most motile cilia of patients with *RPGR* variants were static or showed restricted beat (7/9), while for some cases (#15, 11), a mixture of slow and fast beat cilia was observed (Fig. 2C). For patients with variants in the *RPGR^ORF15^*, in most cases (8/9), cilia beat frequency decreased significantly (Fig. 2C). For cilia beat waveform, besides the normal cilia beat, four different types of beat phenotypes could be identified: 1) static cilia; 2) restricted cilia beat; 3) large cilia beat amplitude with a loss of power and recovery stroke pattern; 4) cilia beat in rotationary and/or uncoordinated manner (Fig. 2D). We quantified the percentage of cells with different cilia beat modes. While most cells with *RPGR* variants (7/9) had static cilia or restricted cilia beat, cells with *RPGR*^ORF15^ variants showed better cilia beat (8/9, 1 has normal cilia beat) but they either lost the waveform or coordination (Fig. 2E).

**Fig. 2.**
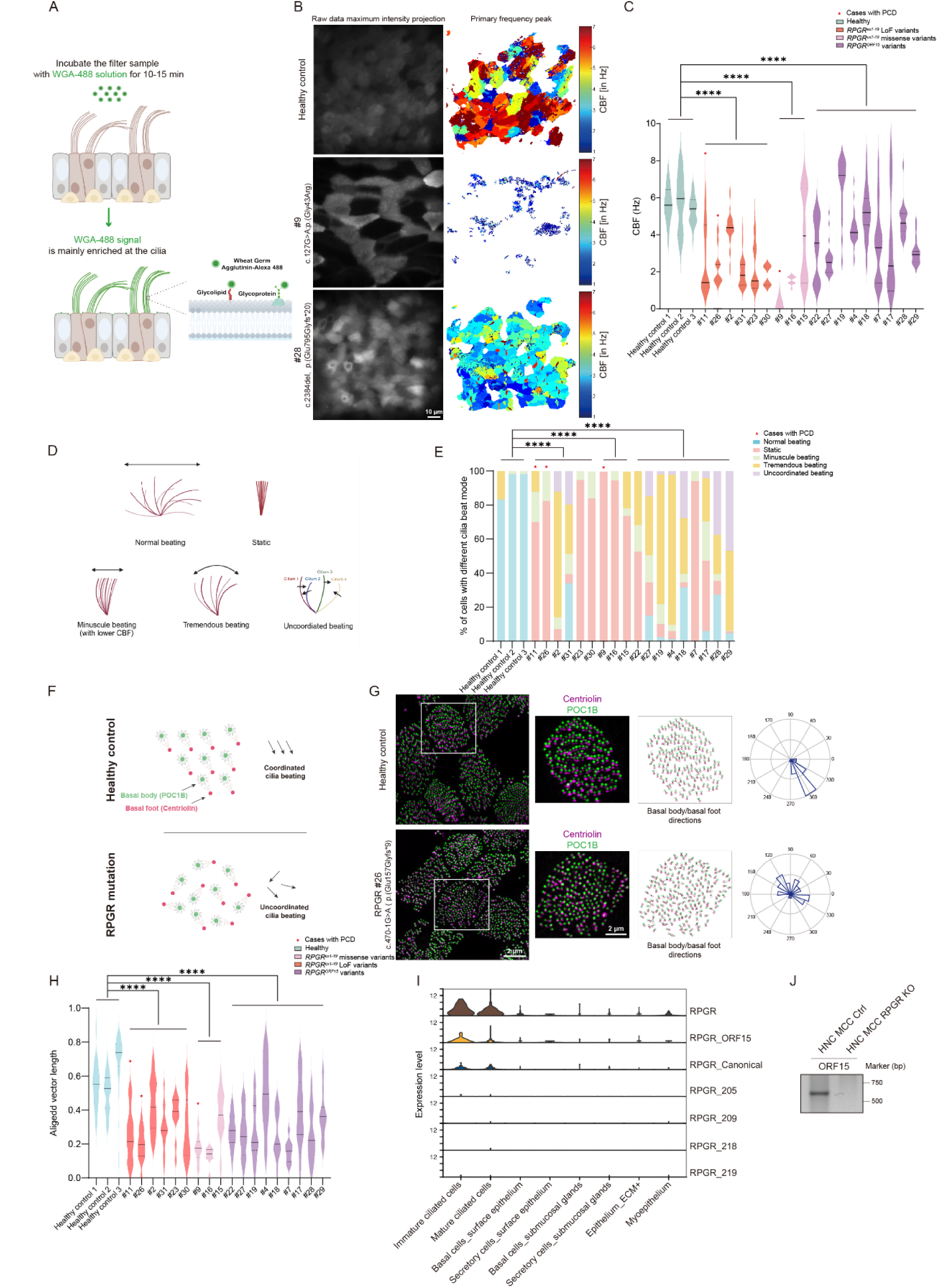
Multiciliated Cells (MCCs) with pathological *RPGR* variants showed impaired and uncoordinated cilia beat. **(A)** A cartoon showing labelling motile cilia with WGA-Alexa 488. **(B)** Pixel frequency map shows cilia beat impairment in the MCC with *RPGR* pathological variants. Scale bar, 10 μm. **(C)** Characterization of cilia beat frequency for all RP patients in this study. n>15 cells per sample. **(D)** Different cilia beat modes identified in *RPGR* loss of function MCCs. **(E)** Characterization of cilia beat waveform distributions for all RP patients in this study. n>16 cells per sample. **(F)** A cartoon showing labeling Poc1B and Centriolin to assess rotational polarity and its disruption in MCCs bearing *RPGR* pathological variants. **(G)** Immunostaining shows rotational polarity in healthy control cells and its disruption for one patient with pathological *RPGR* mutation. **(H)** Characterization of rotational polarity for all RP patients in this study. n>9 cells per sample. **(I)** Single-cell RNA seq revealed the expression of *RPGR* mRNAs and its different isoforms in airway epithelial cells. **(J)** RT-PCR showed the expression of the ORF15 isoform in human airway epithelial cells (HNC cultured at ALI for > 8 weeks). All data are presented as average ± SEM. The center, upper and lower lines represent the median, upper, and lower quartiles, respectively (C, H). n.s. no significance, *, p < 0.05, **, p < 0.01, ***, p < 0.001, ****, p < 0.0001 by two-tailed t-test (C, E, H).

To quantitatively compare beat coordination, we used a 3D-SIM based method we previously developed to examine the rotational polarity of the basal foot (Fig. 2F and 2G), an appendage structure localized at the base of each cilium (*43–46*). In healthy MCCs, all basal feet point towards the direction of the cilia beat, a phenomenon called rotational polarity (Fig. 2F), which is disrupted in PCD patients because of defective cilia beat (*46*). The rotational polarity was disrupted in both cells with *RPGR* (9/9) and *RPGR^ORF15^*(9/9) variants (Fig. 2H).

Taken together, damaging variants in *RPGR* and *RPGR^ORF15^* affect airway MCC cilia beat frequency (9/9, 8/9 for *RPGR* and *RPGR^ORF15^* respectively), waveform (9/9, 9/9), and coordination (9/9, 9/9).

### *RPGR*-RP patients with severe PCD symptoms demonstrate severe cilia defects

Patients with severe clinical respiratory PCD symptoms tend to have *RPGR* loss of function variants or in frame deletions and demonstrate more severe cilia defects. Clinical features of cases # 9, 10, 11, 20, 25, and 26 met the diagnostic criteria of PCD, and all showed shortened nasal cilia length (Fig. 1E). We successfully re-differentiated cells from cases # 9, 11, and 26, and found all showed reduced cilia length and ciliation except for the ciliation of differentiated MCCs from case #11 was not affected (Fig. 1F and 1I). Cilia beat analysis of MCCs from those three cases showed that most MCCs presented static cilia or restricted cilia beat, and the rotational polarity was disrupted in all (Fig. 2C, 2E, and 2H, Table S2).

Despite cases #10& 11 (relatives), 23& 24 (relatives) bearing the same variant leading to alternative splicing (c.934G>T; p.(Glu260_Thr311del)), cases #10, 11 presented with a clinical PCD phenotype and shorter nasal cilia length, while cases #23 & 24 had shorter cilia but longer than #10,11, in addition to milder respiratory symptoms (Fig. 1E). This suggests multifactorial influences in the development of PCD. However, the ALI cells from case #23 showed all cilia phenotypes and he had abnormal respiratory features and hearing loss (Table S2), suggesting this variant does predispose him to motile cilia dysfunction.

Though most *RPGR* RP patients showed motile cilia defects, only a population presented with clearly an abnormal respiratory phenotype, and 6 with a strong PCD phenotype (6/32), all had variants affecting both isoforms. However, in some cases of patients with *RPGR^ORF15^* variants, abnormal cilia, and abnormal respiratory phenotype were observed, suggesting these patients are not immune to respiratory illnesses caused by cilia defects (Fig.1 and 2, and Table S2). Our data provides insight into which RP patients have a high risk of PCD or any respiratory illnesses.

### *RPGR^ORF15^* isoform is expressed in airway MCCs

More severe motile cilia defects are seen in patients with variants that disrupt both isoforms (Fig. 1 and Fig. 2). Though the *RPGR* ^ex1-19^ isoform was once regarded as the sole isoform in airway MCCs (*17*), we first showed that cells from patients with *RPGR^ORF15^* variants also present with cilia defects (Fig. 1E, 1F, and 1I), though the respiratory phenotypes were less severe (Table. S2). To verify if the *RPGR^ORF15^* isoform is expressed in MCCs, we first examined the single-cell sequencing data of fetus trachea generated in a previous study by He, et al (*47*). All *RPGR* isoforms were first treated as a single gene. At the mRNA level, compared to other cell types, *RPGR* was mainly expressed in the MCC progenitors (Foxn4 expressing) and mature MCCs (Fig. 2I). We next separated different isoforms and performed isoform-specific expression analysis. The result for the first time showed that the expression levels of the *RPGR^ORF15^* isoform were comparable to the most expressed *RPGR^ex1-19^* isoform, now confirming that *RPGR^ORF15^* can be expressed in airway cells. The expression of the *RPGR^ORF15^* isoform was upregulated during the early stage of multiciliogenesis (Fig. 2I), indicating its role in cilia formation. To further examine the expression of *RPGR^ORF15^* isoform in MCCs, we performed RT-PCR of the total mRNA transcripts extracted from healthy controls and *RPGR* KO HNCs. The results further support that *RPGR^ORF15^* is expressed in airway cells (Fig. 2J and S4), consolidating the single-cell sequencing data.

### *RPGR* KO MCCs present sparse short motile cilia

To determine the *RPGR* specificity of our findings in patient cells, we generated *RPGR* KO MCCs by CRISPR-mediated genomic perturbation. In brief, either healthy human nasal or bronchial basal cells were edited through infection of lentivirus expressing Cas9 and *RPGR* gRNAs, selected by puromycin, and differentiated at the air-liquid interface into airway MCCs (Fig. 3A). We optimized the protocol by adding ROCK inhibitor Y27632 (*48, 49*), and irradiated fibroblast cells (*50–52*), known to enhance basal cell growth while maintaining their differentiation potential. We adopted two gRNAs reported by Zhang, et al. (Fig. 3B) (*53*), and a combination of these two gRNAs targets exons 2-4 and disrupts the RCC1 domain of both isoforms (Fig. 3C). Using this optimized protocol, we successfully generated *RPGR* KO MCCs evidenced by DNA gel electrophoresis and Sanger sequencing: the percentage of cells in which dual gRNAs worked reached 92.3% and the total KO efficiency was ∼99% (Fig. 3C and 3D). Immunostaining showed a complete absence of RPGR compared to the control (Fig. 3D). We further characterized the cilia properties for all 5 biological replicates, including 2 human nasal cell (HNC) and 3 human bronchial epithelial cell (HBEC) samples, showing that cilia length was affected in all (Fig. 3E and 3F). Quantification of the ciliation level of MCCs showed that, while basal body number was unaffected, ciliation was significantly decreased (Fig. 3E, 3G, and S2C), which suggests that RPGR regulates ciliation but not basal body amplification or docking.

**Fig. 3.**
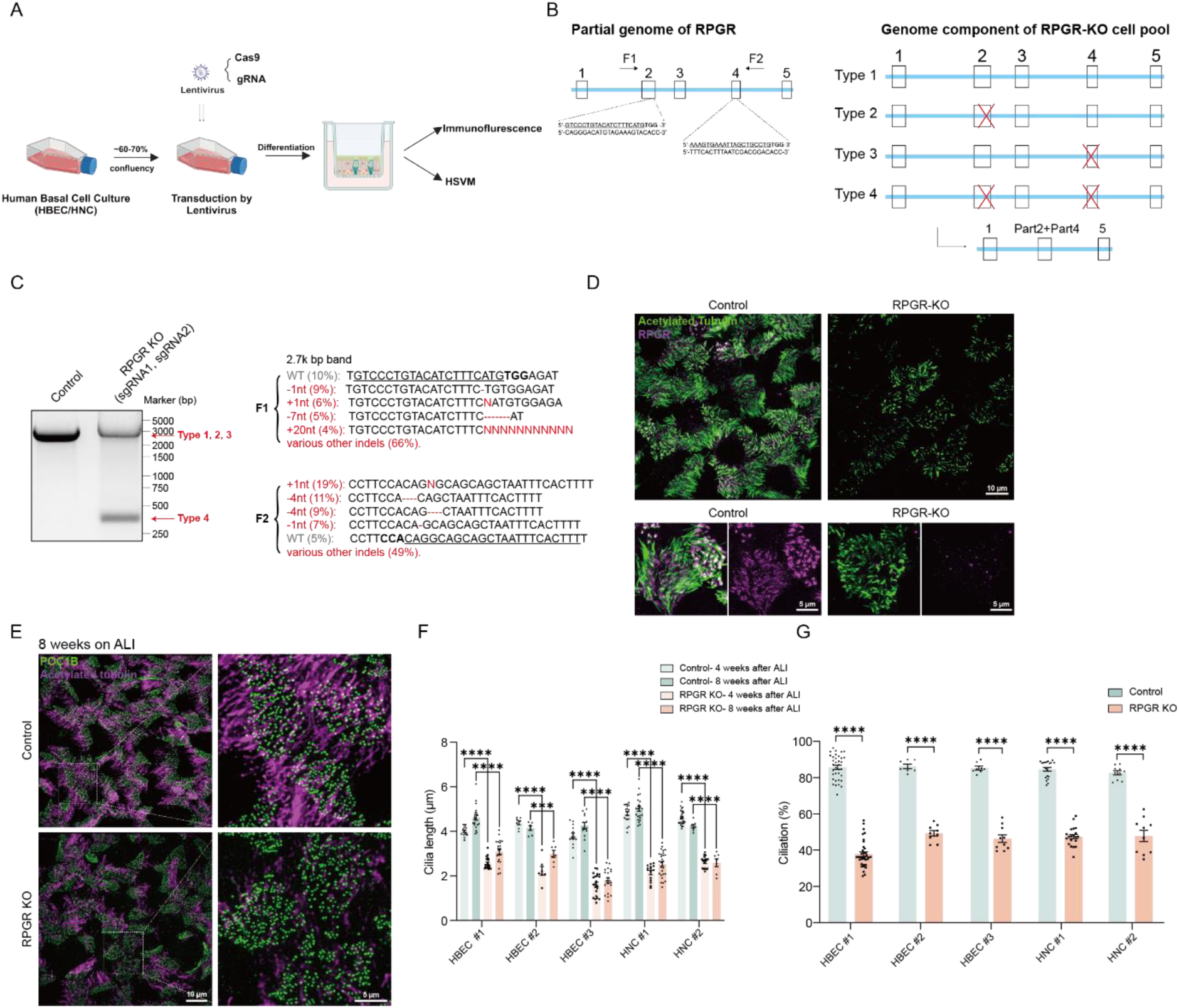
*RPGR* Knockout(KO) Multiciliated cells (MCCs) presented sparse and short motile cilia. **(A)** Workflow of generating *RPGR* KO MCCs. **(B)** The location of guide RNAs and the expected genome edits. **(C)** DNA gel and Sanger sequencing showed high efficiency of *RPGR* KO in MCCs. **(D)** Immunostaining of MCCs with RPGR antibody showed complete RPGR removal in *RPGR* CRISPR KO MCCs. **(E)** Immunostaining with POC1B and acetylated tubulin antibodies showed the reduction of ciliation length and ciliation for *RPGR* KO HBEC/HNC cells. Scale bar 5 μm. **(F)** Cilia length was significantly disrupted in all 5 biological replicates. n>7 cells per sample. **(G)** Ciliation was severely affected for all 5 *RPGR* KO biological replicates. n>7 cells per sample. All data are presented as average ± SEM. n.s. no significance, *, p < 0.05, **, p < 0.01, ***, p < 0.001, ****, p < 0.0001 by two-way repeated ANOVA followed by Sidak’s post hoc test (F, G).

Taken altogether, *RPGR* CRISPR KO airway cells had shortened cilia and reduced ciliation levels, recapitulating the cellular phenotypes seen in the patients.

### *RPGR* KO MCCs present static cilia and/or abnormal cilia motility

Live cell imaging of *RPGR* KO MCCs (Fig. 4C) showed the majority of the cells (68%, N=4) had static cilia, in 4 of 5 biological replicates (Fig. 4B upper and middle, Video S2), leading to disrupted mucociliary clearance (Video S3). We identified large heterogeneity within and among different biological replicates (Fig. 4C). A quantification of cilia beat frequency of the MCCs showed results consistent with patient data. Analysis of the cilia beat waveform, in 4 out of 5 replicates, showed the static cilia were predominant while for the 5^th^ replicate, although most cilia could beat with a normal frequency compared to the control (n.s., 2 technical replicates), both the waveform and beat coordination were abnormal (Fig. 4D).

**Fig. 4.**
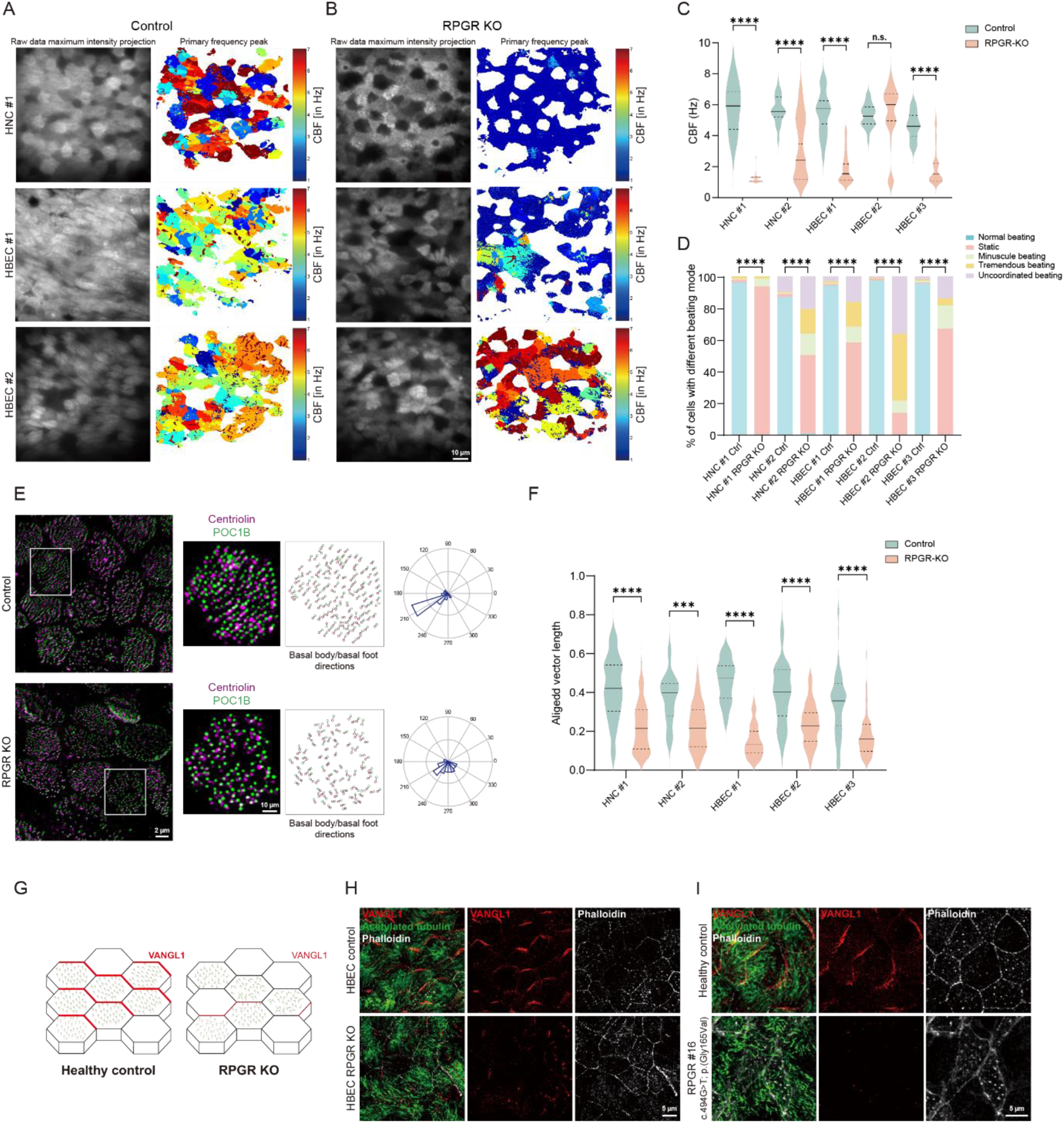
*RPGR* KO MCCs presented with limited motility and defective rotational and planar polarity. **(A, B)** Pixel frequency map shows control MCC cilia beat and its impairment in *RPGR* KO MCCs. **(C, D)** Characterization of cilia beat frequency and beat mode changes for 5 biological replicates in this study. n>27 cells per sample. **(E)** Immunostaining showed rotational polarity in control cell and its disruption for RPGR KO MCCs. **(F)** Rotational polarity was disrupted for all *RPGR* KO samples in this study (N=4). n>13 cells per sample. **(G-I)** Vangl1 mislocalization for MCCs from an HBEC *RPGR* CRISPR KO sample and comparison to an RP patient sample suggesting loss of planar polarity. Scale bar, 5 μm. All data are presented as average ± SEM. The center, upper and lower lines represent the median, upper, and lower quartiles, respectively (C, F). n.s. no significance, *, p < 0.05, **, p < 0.01, ***, p < 0.001, ****, p < 0.0001 by two-tailed t-test (D), or two-way repeated ANOVA followed by Sidak’s post hoc test (C, F).

The rotational polarity assessment of *RPGR* KO cells showed a drastic disruption of beat coordination (Fig. 4E and 4F). As previous studies suggested the disruption of planar cell polarity (PCP) for patient cells bearing *RPGR* variants (*54, 55*), we stained the ALI filters with antibodies targeting the planar polarity protein Vangl1 (*56, 57*). For control cells, Vangl1 was specifically enriched to one side of the MCCs. For either *RPGR* KO HNC, HBEC cells, or patient cells with either *RPGR* or *RPGR^ORF15^* variants, the asymmetrical distribution of Vangl1 was largely lost, also suggesting planar polarity disruption (Fig. 4G-I and S2D).

To summarize, *RPGR* defects caused by CRISPR perturbation lead to either static cilia or cilia with altered motility, disrupting both planar and rotational polarity.

### RPGR locates to the transition zone (TZ) and cilia and its loss does not affect the integrity of the TZ or the localization of axoneme dyneins

To inspect how RPGR regulates motile cilia, we first examined RPGR subcellular localization by 3D-SIM. We stained airway MCCs from controls with a validated RPGR antibody. 3D-SIM provides ∼120 nm resolution (*58–60*) and showed clear transition zone distribution of RPGR by colocalization with transition zone marker RPGRIP1L as well as a weak signal in cilia (Fig. 5A). We further employed STORM (*61–63*), which provides ∼20 nm resolution, to examine the distribution of RPGR and found that it formed a dotted ring pattern at the transition zone for each cilium (Fig. 5B) as well as cilia localized puncta.

**Fig. 5.**
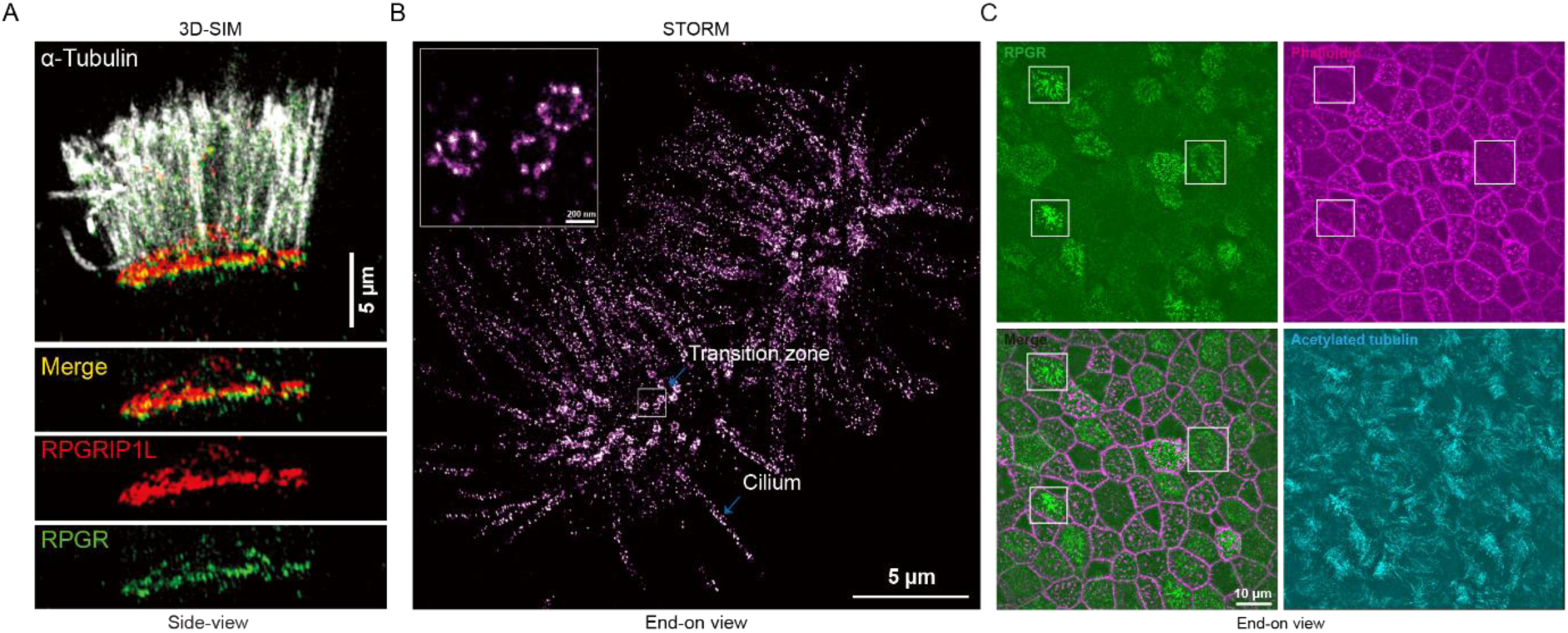
RPGR mainly locates to the transition zone (TZ) and cilia and the exclusive distribution of RPGR and F-actin. **(A)** 3D-SIM showed RPGR mainly locates to the TZ of motile cilia. Scale bar, 5 μm. **(B)** STORM showed RPGR presents dotted ring structure at the transition zone and puncta along cilia. Scale bar, 5 μm and 200 nm for the insert. **(C)** The exclusive distribution of RPGR (green) and F-actin (Phalloidin-Alexa 647 labeled magenta) in the MCCs cultured on ALI for 4 weeks: when RPGR was highly expressed, F-actin signal is weaker and vice versa.

The transition zone (TZ) is a diffusion barrier between the cilia lumen and cytoplasm, allowing selective cargo entry and exit of cilia (*64*). The subcellular localization of RPGR and its interaction with transition zone components suggests that it might be involved in ciliary transportation and cilia composition maintenance. Since most cilia of *RPGR* LOF cells are static, we wondered if the axoneme outer dynein arms (ODA) and inner dynein arms (IDA) are intact. In patient samples with either *RPGR^ORF15^* or *RPGR* variants, ODA marker DNAH5 and IDA marker DNALI1 showed proper localization (Fig. S5A and S5B), consistent with the normal TEM results (Fig. S5C) (*65*). Knowing that RPGR interacts with transition zone components RPGRIP1L, CEP290, and NPHP4 (*66, 67*), we assessed if the loss of RPGR affects the localization of these proteins at the TZ. By staining the *RPGR* KO MCCs with corresponding antibodies, we found that loss of RPGR does not affect the distribution of these transition zone components (Fig. S5D). To assess if RPGR loss affects the organization of TZ, we stained other TZ components such as AHI1, MKS1, MKS3, and CC2D2A and found they were all properly localized (Fig. S5E), suggesting that RPGR is dispensable for the integrity of transition zone.

### *RPGR* KO MCCs form condensed actin meshwork

RPGR has been reported to regulate F-actin polymerization (*68, 69*) and loss of RPGR in hTERT-RPE1 cells induces the formation of actin bundles (*68*). Using an *RPGR* KO mouse model, Roly, *et al.* suggested that RPGR regulates photoreceptor ciliary tip actin dynamics, affecting outer segment membrane turnover (*26, 70*). To investigate if RPGR also regulates actin organization in MCCs, we first looked at the relative distribution of F-actin and RPGR: In ALI 3-week MCCs, when RPGR was highly expressed, Phalloidin-Alexa647 labeled F-actin signal was weaker and vice versa (Fig. 5C). Evaluation of the distribution of F-actin in MCCs using 3D-SIM showed that actin demonstrates dynamic distribution at different stages of multiciliogenesis (Fig. 6A): at ALI 4 weeks, the apical surface actin forms a dense actin meshwork (Fig. 6a left), which is critical for basal body docking (*71, 72*). After ALI 8 weeks, the apical actin meshwork dissolves and the actin patches concentrate at the base of cilia where filamentous F-actin bundles can be readily observed (Fig. 6A right). The filamentous F-actin is supposed to align basal bodies to synchronize cilia beat coordination (*72, 73*). At ALI 8 weeks, 3D-SIM imaging of *RPGR* KO MCCs showed that the apical actin network did not dissolve, which affected the extrusion of the cilia (Fig. 6B). Quantification of the percentage of apical surface covered by F-actin showed that for all 4 biological replicates, the *RPGR* KO cell surface F-actin portion was significantly larger than controls at ALI 8 week (Fig. 6C). To gain further insights into the organization of actin, we applied STORM imaging of Phalloidin-Alexa 647 labelled MCCs and found that for control cells at ALI 8 weeks, the F-actin appeared as bright puncta while for *RPGR* KO MCCs the F-actin appeared as a meshwork covering the apical surface (Fig. 6D).

**Fig. 6.**
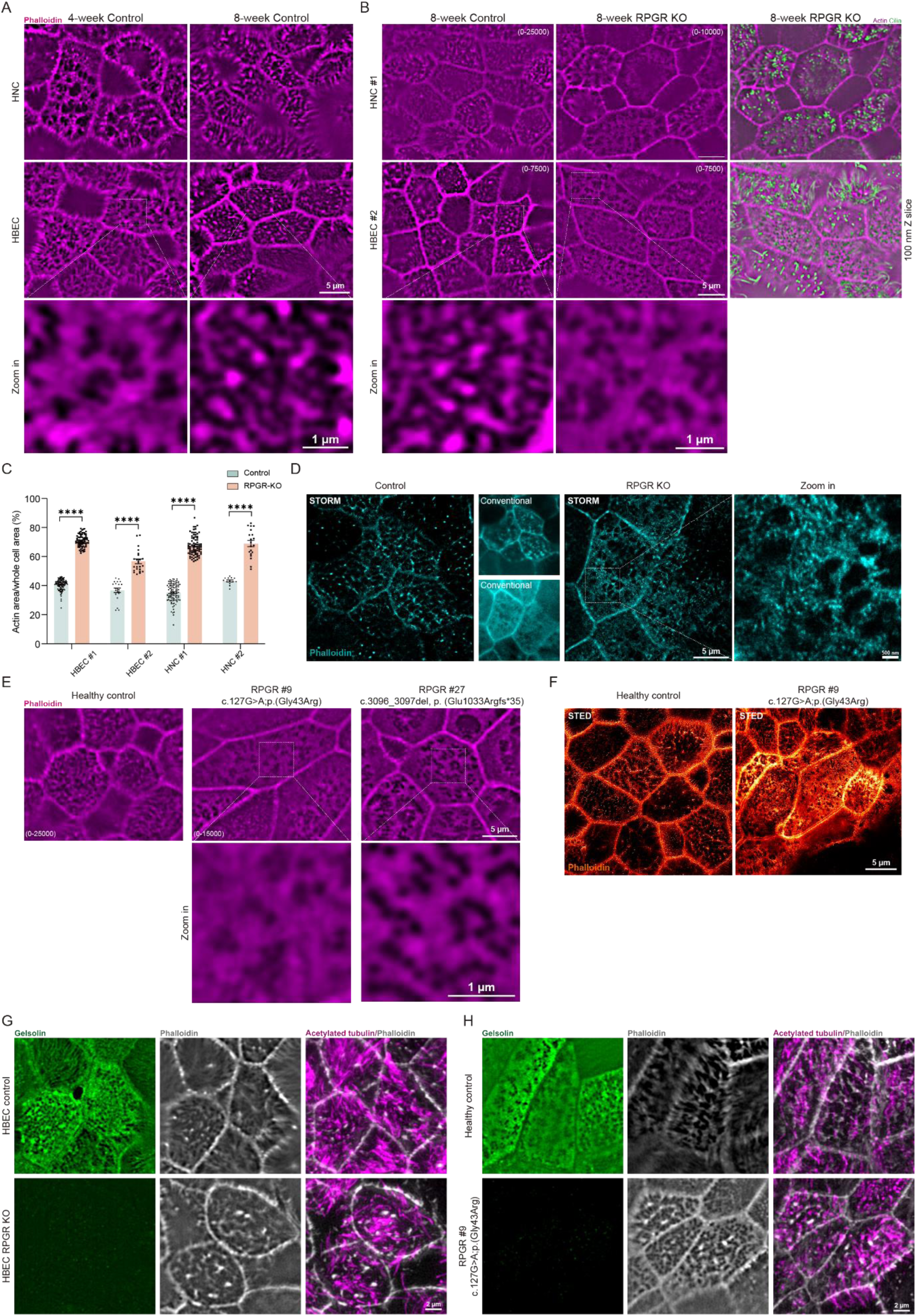
Mature *RPGR* loss of function MCCs present condensed actin meshwork at the apical surface that doesn’t dissolve. **(A)** The different distribution of F-actin at different stages of MCCs: For ALI 4-week samples, F-actin forms dense meshwork at the apical surface; in mature multiciliated cells (8 weeks), F-actin forms puncta at the cilia base connecting into strips, which is important for cilia beat coordination. The zoomed-in image shows the actin meshwork (left) and actin puncta (right) respectively. **(B)** The F-actin meshwork for *RPGR* KO mature MCCs: Instead of forming puncta distributions, F-actin maintains the meshwork that fails to dissolve, similar to 4-week samples. The zoomed-in image shows the actin puncta (left) and actin meshwork (right) respectively. **(C)** The apical surface is covered with more percentage of F-actin in *RPGR* KO HNCs or HBECs. **(D)** STORM imaging of the apical F-actin in both *RPGR* KO cells and healthy control cells. **(E)** The MCCs from patients bearing *RPGR* pathological variants, present with an apical surface that is covered with undissolved F-actin meshwork, similar to *RPGR* KO model. **(F)** STED imaging of the apical F-actin in both RP patient cells with *RPGR* pathological variants and healthy control cells. **G, H)** Apical Gelsolin was diminished in 4-week *RPGR* KO MCCs. **(G)** Apical gelsolin was similarly decreased for *RPGR* KO HBEC MCCs. **(H)** Apical gelsolin was decreased for the MCCs from RP patients with *RPGR* pathological variants. All data are presented as average ± SEM. n.s. no significance, *, p < 0.05, **, p < 0.01, ***, p < 0.001, ****, p < 0.0001 by two-way repeated ANOVA followed by Sidak’s post hoc test (C).

Analysis of MCCs from RP patients showed results mostly consistent with that observed in *RPGR* KO cells. For most patients with loss of function variants in *RPGR* (N=6/10) or *RPGR^ORF15^* (N=6/9), the apical actin meshwork in 8-week MCCs could be readily discerned (Fig. 6E and S5A) and STED −50 nm resolution-imaging of F-actin in patient #9 further supported our findings (Fig 6F). However, in some patients (#2, 4, 11, 19, 29, and 31), the F-actin accumulation seemed to be milder/normal (n=3/10 for *RPGR* variants and n=3/9 for *RPGR^ORF15^ variants*) (Fig. S6B). Cases 4, 19, and 29 are *RPGR^ORF15^* variants, and # 19 and # 29 had normal lung assessments. Case 11 (*RPGR*) was 14 years of age at the time of assessments, though he had PCD, the ocular phenotype was not advanced, as expected for that age. However, Case 2 (*RPGR)* only had a mild airway anomaly assessment, a milder eye phenotype than expected for his age, and showed no F-actin changes. This patient retained decent visual acuity (20/60, Table S2) for 36 years of age. Case 31(*RPGR*) had a normal lung assessment. Of the cilia phenotypes studied, 12 of 19 patients had F-actin changes (Table S3) while this involved (6/9) of those with ORF15 variants. The variability observed is clearly not solely genetically driven as frameshift variants lead to milder phenotypes (Case #2) and siblings of similar age have different phenotype severity (Cases #23 and #24).

In photoreceptor and hTERT-RPE1 cells, RPGR regulates actin dynamics through interaction with Gelsolin and Cofilin (*26, 70*). Staining for Gelsolin for both 4-week control and *RPGR* KO MCCs showed that in both *RPGR* KO HNCs and HBECs (N=5), the cell surface, the layer where F-actin mainly locates, had temporarily diminished Gelsolin signal, suggesting that RPGR might regulate F-actin dynamics through locating Gelsolin (Fig. 6G and S7A). Gelsolin exists in active and non-active forms and it is the active Gelsolin that binds to F-actin (*74, 75*). Reduction of Gelsolin binding to F-actin in *RPGR* KO MCCs suggests that RPGR might be involved in Gelsolin activation. We further examined the distribution of Gelsolin in MCCs bearing different *RPGR* variants (Cases #9, #11, #15, #16, #23, and #30, 6/6) and found a consistent decrease of the Gelsolin that colocalized with the surface F-actin (Fig. 6H and S7B).

In summary, RPGR regulates Gelsolin distribution spatiotemporally, and *RPGR* KO MCCs and MCCs with *RPGR* pathological variants maintained an actin meshwork that did not dissolve (Fig. 7I).

**Fig. 7.**
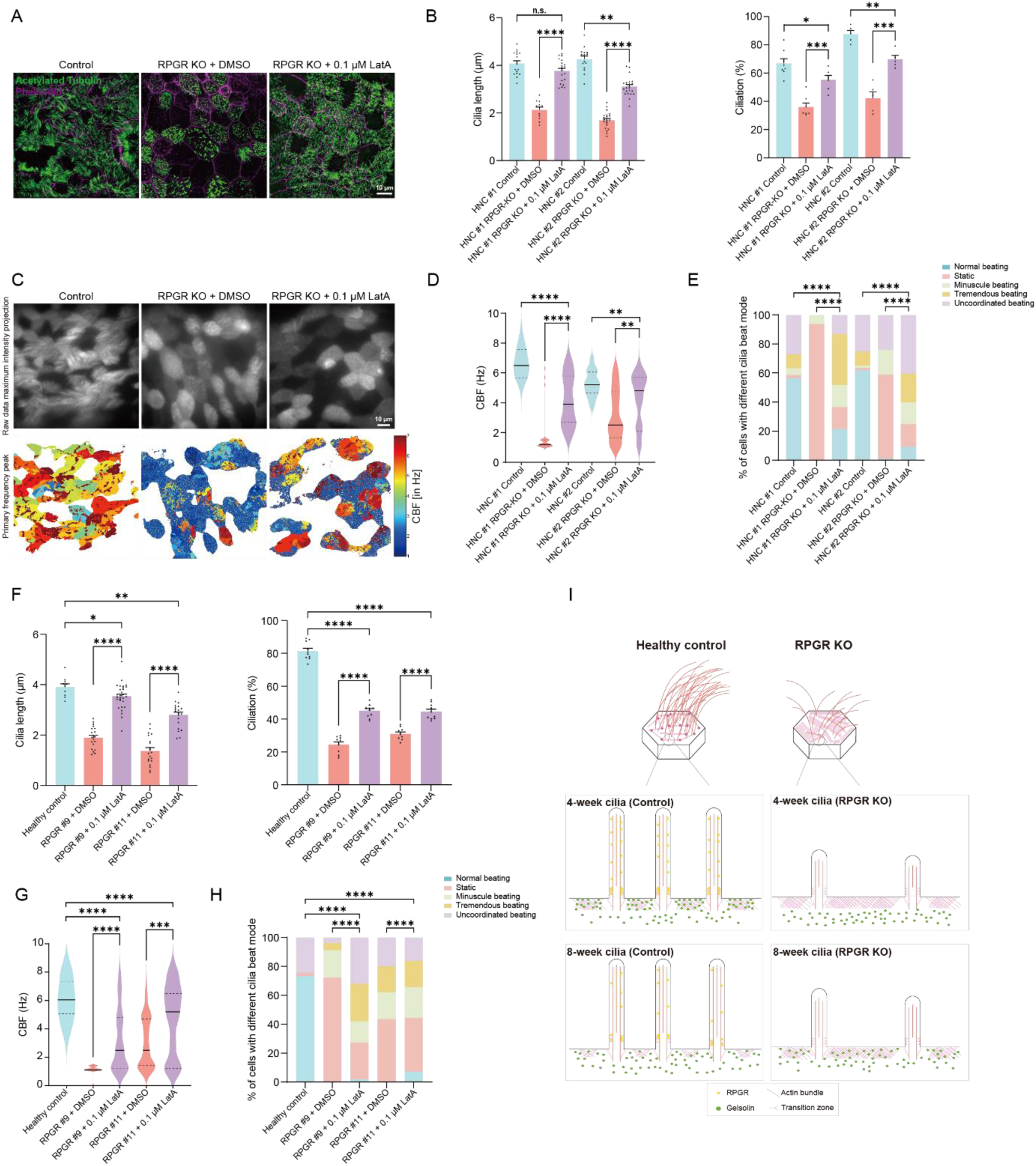
Latrunculin A (LatA) treatment ameliorated the motile cilia defect caused by RPGR loss of function in both *RPGR* KO model and RP patients MCCs. **(A, B)** The ciliation and cilia length anomalies in *RPGR* KO MCCs were ameliorated by LatA treatment. (A) Cilia were stained with acetylated antibodies (green) and the cell boundary was labelled with Phalloidin (magenta). The images showed the reduction of ciliation and cilia length in *RPGR* KO MCCs (middle) and the rescued phenotypes after LatA treatment (right); (B) Image quantification showed the improvement of cilia length and ciliation for 2 *RPGR* KO biological replicates; **(C-E)** The HNC cilia beat anomalies in RPGR loss of function MCCs were partially rescued by LatA treatment. (C) Cilia were stained with WGA-488 (upper) for cilia beat observation and the lower panel is the corresponding pixel signal fluctuation frequencies. The images showed a reduction in cilia beat in *RPGR* KO MCCs (middle) and increased frequency after LatA treatment (right). (D) Quantitative analysis showed the cilia frequency changed before and after LatA treatment. (E) Cilia beat waveform was improved after LatA treatment; **(F-H)** The ciliation, cilia length, and cilia beat properties in MCCs from patients with *RPGR* pathological variants (#9: c.127G>A, p.(Gly43Arg), #11: c.934G>T,p.(Glu260_Thr311del)) was alleviated by LatA treatment. (F) Cilia length (left) and ciliation level (right), (G) Cilia beat frequency, (H) Cilia waveform; **(I)** The proposed model of this study: RPGR regulates F-actin through temporarily recruiting/activating Gelsolin. Without RPGR, F-actin meshwork accumulates at the MCC apical surface, preventing ciliation, cilia elongation, and cilia beat is severely impaired. All data are presented as average ± SEM. The center, upper, and lower lines represent the median, upper, and lower quartiles, respectively (F, I). n.s., no significance, *, p < 0.05, **, p < 0.01, ***, p < 0.001, ****, p < 0.0001 by two-tailed t-test (B, D, F, G, H, I, J).

### Motile cilia anomalies caused by *RPGR* variants are improved with Latrunculin A

If RPGR truly regulates motile cilia through F-actin, treatment with actin polymerization inhibitors such as Latrunculin A (LatA) (*74, 76*) should rescue or ameliorate the phenotype. Because previous studies showed that overexpression of active Gelsolin can rescue the ciliation issue in *RPGR* KD hTERT-RPE1 cells (*70*), we first worked on hTERT-RPE1 for its short turnaround time. From our *RPGR* KO hTERT-RPE1 cell pool, we found the ciliation and cilia length defects were consistent with previous studies (*53*) and our motile cilia data (Fig. S8A,B and C). Treatment of hTERT-RPE1 control and *RPGR* KO cells with 0.2 μM LatA for 24 hours led to 47.4 ± 2.0 (n=3) % ciliation of control cells and 50.2 ± 3.6 (n=3) % ciliation of the *RPGR* KO cells (Fig. S8D and E). The cilia length of control cells reached 4.3 ± 0.4 μm (n=3) while the *RPGR* KO cells had a mean cilia length of 3.9 ± 0.3 μm (n=3) (Fig. S8D and S8E), which was not significant compared to controls. This suggests that LatA can rescue the cilia phenotype caused by loss of RPGR and that RPGR works upstream of actin in cilia regulation. We also applied LatA on the patient-derived fibroblast cells from RP patients #9 and #12 that have cilia defects (Fig. S8F and S8G) and found that LatA treatment partially rescued the phenotypes (Fig. S8H and S8I).

To investigate if LatA treatment could rescue motile cilia, we applied 0.1 μM LatA on ALI cultured *RPGR* KO MCCs starting from the 2^nd^ week and observed cilia properties at the 5^th^ week as RPGR strongly expresses during the early stage (Fig. 2I, S9A, S9B, and S9C). The cilia length and ciliation were partially rescued (N=2, Fig. 7A and 7B). Since F-actin is also required for basal body alignment, we reason that longtime LatA treatment might also affect cilia beat, we thus evaluated the cilia beat after 2-week LatA treatment. For *RPGR* KO cells, which most had static cilia, showed that after LatA treatment the percentage of cells with static cilia decreased and the percentage of cells with normal or rigid cilia beat increased significantly (N=2, Fig. 7C-E, Video S4), suggesting that actin depolymerization could indeed ameliorate the phenotypes caused by RPGR defect. The evaluation of LatA treatment on patient cells bearing pathological *RPGR* variants (#9 and 11, with PCD) showed results consistent with the improvement observed in *RPGR* KO cells (Fig. 7F-H, Video S5).

In summary, the structural anomalies and motile cilia dysfunctions caused by RPGR variants can be partially rescued by LatA treatment.

## DISCUSSION

In this study, we investigated the mechanisms by which *RPGR* defects affect motile cilia functions. By characterizing the cilia from 28 patients with a wide range of *RPGR* variants and that of *RPGR* KO MCCs using advanced microscopy, we first reveal that RPGR regulates ciliation, cilia length, and cilia beat properties in airway MCCs. Our results suggest that these changes might be caused by the reduction of active Gelsolin at the apical surface, resulting in the failure of actin meshwork clearance. Treatment with LatA ameliorated all cilia phenotypes (Fig. 7). This is the first mechanistic study of RPGR’s role in cilia motility. It reveals a new mechanism for PCD and complements current studies of RPGR in photoreceptors and hTERT-RPE1 cells.

PCD remains an underdiagnosed disorder of the motile cilia which affects airway health outcome(*77, 78*). A delayed diagnosis can lead to patients’ lungs deteriorating, even to lung transplantation or death (*8*). Patients with *RPGR* variants are usually referred to eye clinics because they have RP while their respiratory symptoms can be mild and undiagnosed. *RPGR* has long been recognized as a PCD causative gene, but the diagnosis of *RPGR*-PCD can be challenging and certainly under-recognized as the cilia phenotype can be different than other causes of PCD. PCD caused by defects in other proteins may be diagnosed by demonstrating classic cilia ultrastructural defects by transmission electron microscopy (TEM), such as missing outer and/or inner dynein arms. However, *RPGR* defects do not result in classic ciliary ultrastructural defects (*31*), therefore in these cases, TEM is not useful in diagnosing PCD in RP patients. Undiagnosed PCD patients were identified through this study. This work clearly highlights how *RPGR* variants affect various cilia properties, which could possibly be applied for *RPGR*-PCD diagnosis. We showed that cilia length, motility, and rotational polarity were abnormal even when the patient’s respiratory phenotype was mild or normal.

Importantly, this study shows how different *RPGR* variants contribute to impaired mucociliary clearance, as well as airway pathology. The variants affecting both isoforms appear to be associated with a stronger clinical phenotype however we have shown that all *RPGR* variants affect cilia health. There was variability in clinical outcomes even in patients carrying the same *RPGR* variant. This could possibly be explained by different environmental factors or genetic backgrounds and interactions. Unlike the early studies suggesting that *RPGR^ex1-19^* was the sole isoform in airway MCCs, we showed that the *RPGR^ORF15^* isoform is expressed in MCCs and that variants in *RPGR^ORF15^* could affect cilia health, explaining the respiratory signs and symptoms observed in some *RPGR^ORF15^*variant carriers.

We found that RPGR in MCCs localizes both to TZ and cilia. Since the C terminal of the canonical RPGR is prenylated (*69*), the cilia distribution of RPGR should be associated with the cilia membrane, which is different from most PCD causative genes. Consistently, we did not identify changes in DNAH5 and DNALI1 distributions. Surprisingly, we did not find localization changes for either RPGR interacting proteins or transition zone components.

Our work revealed that actin turnover appears to play a central role in the regulation of motile cilia by RPGR (Fig. 7K). In control cells, F-actin first forms an apical meshwork, which is presumably important for basal body docking and thus ciliation. The apical meshwork later dissembles and forms actin paths under basal bodies or bundles aligning basal bodies for cilia stabilization and beat synchronization. Though previous studies in *Xenopus* suggested a two-layer of actin existing on the surface of the MCCs (*76, 79*), we only identified one, which may reflect organism differences, or that our staining lacks a good basal body reference marker. Unlike control cells, mature *RPGR* LOF MCCs retained the condensed meshwork which restricts ciliation and cilia elongation, and limits cilia beat. This phenotype likely relates to the reduced level of active Gelsolin observed at the apical surface, which is most obvious for 4-week MCCs. Cofilin is also involved in the regulation of F-actin by RPGR (*70*). We could not explore its role further due to the poor performance of the antibody. Treatment with LatA partially rescued ciliation and cilia length and improved the cilia beat defect. Because of the toxicity of LatA, we were not able to further enhance the LatA treatment. Phalloidin staining of the F-actin suggests although the actin meshwork signal was weakened after LatA treatment, the actin meshwork still existed (data not shown). Another possibility is that mature MCCs also need actin bundles for cilia beat. LatA treatment might prevent the actin bundle formation so full rescue of cilia beat may not be possible.

In photoreceptor connecting cilia, RPGR regulates membrane turnover through actin, key to the stability of this transport system (*26, 70*). Here we show that RPGR also relies on actin to regulate motile cilia in MCCs. Our study reveals that RPGR utilizes the same pathway to realize different outcomes in different cell types.

Disruption of rotational but not planar polarity is a common feature of PCD and has been proposed to aid PCD diagnosis (*46, 80, 81*). Our study here finds that loss of RPGR affects both planar and rotational polarity. RPGR has been reported to be associated with the PCP pathway by affecting the Dvl through the proteasome in hTERT-RPE1 cells (*55*). This is the first study showing the loss of Vangl1 at the cell junctions. Since the downstream of the PCP pathway involves actin dynamics, we speculate that the PCP pathway might also be involved in the regulation of actin polymerization by RPGR. However, it is unclear how RPGR regulates the distribution/expression of Vangl1. Because of the lack of other reliable PCP antibodies, we did not further explore this question.

The study has several limitations in addition to patient recruitment for a rare disease during the COVID-19 pandemic: heterogeneous *RPGR* KO cell pools were generated rather than a homogenous cell line, which could contribute to the heterogeneity of the cilia beat in *RPGR* KO MCCs; Our study was also limited to human cell ex vivo models and it needs further in vivo investigation for mechanistic studies such as using mouse model, however, the mouse PCD phenotype is usually mild regarding respiratory symptoms (*80, 82–84*); Although we identified a role of F-actin turnover in the pathogenesis of *RPGR*-PCD, we cannot exclude the role of other interacting pathways in the alteration of motile cilia phenotypes. It might be worthwhile to assess motile cilia composition changes using systematic ways such as proteomics or cryoEM (*85–89*). RPGR localizes at the TZ and cilia, but the actin meshwork forms beneath the apical cell membrane. How RPGR regulates actin dynamics spatiotemporally is worth further investigation.

In summary, our work showed for the first time that *RPGR*^ORF15^ is expressed in airway MCCs, and variants in both isoforms could lead to motile cilia defects in symptomatic and asymptomatic patients. Variants in *RPGR* led to a more severe airway and cilia phenotype but only a few developed PCD. As this study was not longitudinal, we do not know about the progression of these observations. This study reveals the actin pathway is involved and could be manipulated to treat *RPGR*-PCD and opens new avenues of research for *RPGR*-related diseases and opportunities for improved diagnosis and patient management.

## Materials and methods

### Study design

The purpose of this project is to study the mechanism of PCD caused by RPGR defects. Thirty-two RP patients were recruited and most showed different respiratory symptoms. *RPGR* CRISPR KO multiciliated cells were also generated to validate the cellular phenotypes observed in patient cells. Super-resolution microscopy, live cilia beat observation and drug-related rescue experiments were carried out to characterize the cellular phenotypes and to reveal the mechanism.

### Patient Phenotype Characterisation

All patient participants had a comprehensive eye exam in addition to optical coherence tomography (OCT) imaging, kinetic visual field (Goldmann), and electroretinography when possible (Table S2). The respiratory phenotypes were characterized using PCD-specific protocols (questionnaire, natural history, nasal nitric oxide measurements (nNO), chest X-ray, spirometry, and physical exam) (Table S2).

### Cell collection and culturing

Human nasal cells (HNC) were obtained from six healthy volunteers at HKUST and the Hospital for Sick Children (REB# HREP-2021-0268 (HKUST), 1000005895(SickKids)), and from patients with *RPGR* variants at the Hospital for Sick Children using cytology brushes as previously described (*46, 90*). For all the measurements, the researchers were blind to the genetic information of the subjects. After nasal brushing, HNCs were collected in the pre-warmed BEBM medium (Lonza, cc-3170), washed, and subcultured until confluence. Cells were subsequently seeded on 24-transwell inserts and differentiated with PneumaCult-ALI media (Stem Cell Technologies, 05001) following the manufacturer’s protocol. HBECs were sourced from Lonza (cc2504S) and expanded and differentiated following the manufacturer’s protocol. The BEBM and PneumaCult-ALI media were supplemented with 100 μg/mL Vancomycin, 80 μg/mL Tobramycin, 50 μg/mL Gentamicin, and 1X Antibiotic-Antimycotic. HEK293T cells (gift from Ting XIE, HKUST) were cultured with high-glucose DMEM media (ThermoFisher Scientific, 11965084) supplemented with 10% fetal bovine serum and 1% streptomycin/penicillin. hTERT-RPE1 (ATCC CRL-4000) cells were cultured with DMEM/F12 media (ThermoFisher Scientific, 11320033), supplemented with 10% fetal bovine serum, 1% streptomycin/penicillin and 0.1 mg/mL hygromycin. Fetal bovine serum was removed from the medium to induce ciliation of hTERT-RPE1. Fibroblast cell lines were established at Biobank (The Hospital for Sick Children) as described. Briefly, skin biopsies of the upper arm of patients and controls were collected in sterile Alpha MEM medium (AMEM, Wisent, #310-022-CL) with 15 % fetal calf serum, and 1% streptomycin/penicillin. In a 60mm dish, biopsies were cut into small pieces and collagenase was added. Each coverslip was added to the top of each skin piece, so they keep contact with the top of the pieces to grow. The same supplemented AMEM was added and samples were incubated (37°C and 5% CO_2_). After 1.45 hours, pieces and media with collagenase were transferred and centrifuged. The pellet was pipetted vigorously to break up the pieces and seeded on a T25 flask with the AMEM media supplemented with 15% fetal calf serum for another 3 continues days. For further expansion, purified skin fibroblasts from RP patients were cultured in MEM alpha (ThermoFisher Scientific, 12561056) with 10% fetal bovine serum. The growth medium was changed with Opti-MEM (ThermoFisher Scientific, 31985070) to induce the ciliation of skin fibroblasts.

### Plasmid construction and lentivirus preparation

The lentiCRISPRv2 (Addgene#52961) plasmid was digested with BsmBI-v2 (NEB, R0739), and then ligated with annealed oligos. The guide sequences for *RPGR* KO are as follows (*53*): Guide 1: GTCCCTGTACATCTTTCATG and guide 2: AAAGTGAAATTAGCTGCCTG. For lentivirus packaging, HEK293T cells were cultured with high-glucose, sodium pyruvate-supplemented DMEM media (ThermoFisher Scientific, 11995073) with 10% fetal bovine serum, and transfected with pMD2.G (Addgene #12259), psPAX2 (Addgene #12260), and the lentiviral gRNA plasmid with the 1:10:10 mass ratio by applying PEI MAX (Polysciences, 24765) in Opti-MEM. The medium was changed the next day, and the viral supernatant was harvested 48h post-transfection, filtered with 0.45 μm filter, concentrated with PEG-it virus precipitation solution (ExCell Bio, LV810A-1), and frozen in −80 °C freezer for later use.

### Generation of CRISPR KO cells

We generated *RPGR* KO pools in hTERT-RPE1 cells using lentivirus transduction at a MOI of 2 (each cell receives a copy of lentivirus expressing gRNA1 and a copy of lentivirus expressing gRNA2). The resultant KO pool was selected with puromycin treatment for 7 days. The whole genome DNA was extracted, and the KO region was amplified using designed primers (Forward: ACAAGGGGTTTGTATGGATAAA and Reverse: AAAACCCTAATTTTACTGTTGCC). The KO efficiency was evaluated by DNA gel electrophoresis and by Sanger sequencing, which was analyzed by the ICE CRISPR analysis tool (SYNTHEGO). HNC/HBEC *RPGR* KO pools were generated using lentivirus transduction. Low passage basal cells (P01-P02) were cultured on collagen (STEMCELL, 07001) coated 6-well plate in BEBM medium, and incubated until cells reached 30-50% confluent. For lentivirus transduction, the BEBM medium was supplemented with 15 μg/mL polybrene (Santa Cruz, sc-134220) and 20 μmol/mL HEPES (ThermoFisher Scientific, 15630080), and the lentivirus transduction was conducted at a MOI of 2. When basal cells reached ∼80% confluent, they were sub-cultured with feeder cells in DMEM medium, supplemented with 7% fetal bovine serum, 30% Ham’s F-12 nutrient mix (ThermoFisher Scientific, 11765047), 24 mg/L adenine (Sigma, 1152), 4.5 mg/L hTGF (ThermoFisher Scientific, AF-100-15), 1 μg/L hydrocortisone (StemCell, 7925), 10 mg/L insulin (Sigma, I9278), and 8.6 μg/L cholera toxin (Sigma, C8052). The feeder cells for co-culturing were generated by irradiation of puromycin-resistant NIH-3T3 cells (3000-5000 rads of gamma radiation). The resultant KO pool was selected in 1 μg/mL puromycin for 7-10 days until basal cells reached confluency, and seeded on 24-transwell inserts with a density of ∼5.0 x 10^5^ cells/cm^2^. When cells reached confluency, the medium was changed to PneumaCult-ALI for differentiation. The genomic DNA was extracted from a portion of genomic edited basal cells for KO efficiency evaluation.

### Immunofluorescence

Samples were fixed with 4% paraformaldehyde (Sigma, 158127) and 0.1% glutaraldehyde (Electron Microscopy Sciences, 16019) for 15 minutes, then reduced with 0.1% sodium borohydride (Sigma, 2643) for 6-7 minutes, and blocked with 3% bovine serum albumin (Sigma, A8806) and 0.2% Triton-X 100 in 1xPBS (pH 7.4) for 30 minutes. For methanol fixation, samples were fixed with −20°C histological grade anhydrous methanol for 30 minutes and then blocked with 3% bovine serum albumin and 0.05% Tween 20 in 1xPBS (pH 7.4) for 30 minutes. Optionally, to preserve the morphology of motile cilia, the Transwell filter samples were treated with 0.01% Triton-X 100 in 1xPBS (pH 7.4) for 1 minute before methanol fixation.

Primary antibodies were incubated at 37°C for 3 hours or 4°C overnight. The antibodies used in this study were summarized in Table S4. Secondary antibodies were incubated at 37°C for 30 minutes. DAPI (1 μg/mL) was incubated at 37°C for 1 hour and Phalloidin-Alexa 647 (1: 50) was incubated at 37°C for 7 hours. All washing steps were processed with 1xPBS. Samples were mounted with 0.5% n-propyl gallate and 80% glycerol in 1xPBS, sandwiched between coverslips (25 mm, No. 1.5 H, Deckglaser) and slides, and sealed with nail polish.

### Live cell cilia beat observation

Transwell cultured airway epithelial cells were gently rinsed with PBS to remove mucus and cell debris. The filters were then cut off the Transwell, and incubated with PneumaCult-Ex media containing 100 μg/mL wheat germ agglutinin (WGA)-Alexa 488 (ThermoFisher Scientific, W11261) at 37°C for 20 minutes. The medium was then changed to PneumaCult-Ex media. The labeled filter was put into a confocal dish (MatTek, P35G-1.5-14-C) with the sample side facing the coverslip for imaging. WGA-488 labeled samples were captured at around room temperature (23℃) under an inverted microscope and a 488nm excitation laser was used to excite the fluorophores and the cilia beat data were collected using a 100x/1.50 oil immersion objective lens. Videos were recorded at 50 frames per second (fps) for 10 seconds. For the beads propelling experiment, Transwell cultured 8-week airway epithelial cells were gently rinsed with PBS to remove mucus, and the whole filter was put onto a confocal dish with the sample side facing against the coverslip for imaging. Fluorescent beads with a diameter of 1 μm (ThermoFisher Scientific, F8821) were diluted 1:50 with 1X PBS, and the diluted beads were added to the apical surface. The propelled beads were captured at around room temperature (23℃) under an inverted microscope and a 561nm excitation laser was applied to excite the fluorescent beads, and the data were collected using a 60x/1.20 water immersion objective lens. Videos were recorded at 50 fps for 10 seconds.

### SIM imaging and data analysis

3D-SIM datasets were collected using a Zeiss Elyra 7 with Lattice SIM with alpha Plan-Apochromat 63x/1.46 Oil immersion objective lens and with an additional 1.6x optovar. The fluorophores were excited with 488 nm (500 mW), 561 nm (500 mW), and/or 647 nm lasers (500 mW) tuned to 0.5-10% power through the ZEN software. For each image field, grid excitation patterns were applied on the sample plane and collected for thirteen phases. The fluorescence was collected by the objective and filtered by proper band pass mirrors before reaching the camera. Two PCO edge sCMOS cameras were used to acquire a 5-10 μm thick Z stack with 100 nm per slice. The raw data was reconstructed using the SIM module of ZEN Software. Channel alignment was conducted using a calibrated file generated by Tetraspec beads (ThermoFisher Scientific, T7279). Maximum-intensity projection images were produced for subsequent analysis.

### Cilia length measurement

Side-view multiciliated cells on the maximum intensity projection images were chosen for measurements. The cilia length was obtained by measuring the distance from the cilia tip to the base using the free drawer tool of Zeiss ZEN. The average length of 10 cilia within one cell was defined as the cilia length of each MCC.

### Ciliation level

The ALI filter samples were fixed with methanol and stained as above described with antibodies against acetylated tubulin and POC1B. For each cell, the number of cilia and basal bodies were manually counted using the software Fiji ImageJ, and the ratio of the total number of cilia to the total number of basal bodies was calculated as the ciliation level.

### Rotational polarity analysis

8-week filter samples were fixed with methanol as above described, basal bodies were stained with POC1B antibodies and basal feet were stained with Centriolin antibodies. Samples were observed with 3D-SIM and the maximum intensity projection (MIP) of the reconstructed images was exported as ome.tiff (8 bit) for rotational polarity analysis in MATLAB (*46*). Briefly, binary images were first generated by applying a threshold for two channels. Individual basal body and basal foot were identified in the binary images, paired by pairwise nearest neighbour search, and filtered with a 600 nm cutoff. The basal body-basal foot pairs within one cell were determined by manually defining the contour of the cell with the MATLAB free drawing tool. The direction of each pair was the direction from the center of the basal body to the center of the basal foot. The cilia beat coordination in one cell was represented by the aligned vector length –an average of all basal body basal foot direction within one cell-with value close to 1 meaning full coordination and 0 meaning no coordination.

### STORM imaging and data analysis

For RPGR STORM imaging, cell suspensions were generated by digesting the filter samples with 0.025% Trypsin and resuspending disassociated cells in PneumaCult-Ex media. The cells were then spread and dried on coverslips (25 mm, No. 1.5 H, Deckglaser). The cells were immunostained with primary antibodies against RPGR (1:100 dilution), and then labeled with goat anti rabbit F(ab’)2 Alexa 647 (ThermoFisher Scientific, A21237). For actin STORM data, the filter sample was fixed with PFA and GA, and then labeled with Phalloidin-Alexa 647. The labeled sample was mounted with imaging buffer. The imaging buffer contains 50 mM Tris (pH 8.0), 10 mM NaCl and 10% Glucose, and is supplied with 50 mM β-mercaptoethanol (Sigma, 2661), 56 mg/mL Glucose oxidase (Sigma, 3231), and 17 mg/mL Catalase (Sigma, C9322). Concaved slides were used and an imaging buffer was added between coverslip and slide. The slide was sealed with nail polish. The labeled samples were imaged with an Abbelight SAFe360 setup with 100x/1.50 Oil immersion objective lens, and the 647 excitation laser (500 mW) power was set at 10% to collect the conventional image, and with 100% to switch the fluorophores to dark state and for STORM imaging. The 405 laser (100 mW) power was manually adjusted from 0 to 100% for fluorophores activation. The exposure time was set to 20-50 ms and 60,000∼200,000 frames were collected by a sCMOS Hamamatsu ORCA Fusion camera. Acquired STORM data was then processed by the Abbelight NEO Analysis software for Gaussian fitting. Briefly, after background estimation and removal, the detected molecules were fitted, and the corresponding lateral and axial positions of each activated fluorescent molecule were extracted. Then the super-resolution image was reconstructed based on the coordinates after drift-correction by cross-correlation. The precision was averaged from the localization precisions for all the single molecule events, and the resolution was around 20 nm.

### STED imaging and data analysis

For Phalloidin STED imaging, the Phalloidin-Alexa 647 labeled filter samples were observed using the STEDYCON (Abberior) Super-Resolution Microscope with 100x/1.4 Oil immersion objective lens, and the ultra-short pulse red laser (800 μW) excitation power was set at 1%, and the 775 nm depletion laser (1.2 W) power was set at 20% to collect the STED datasets. Acquired STED data was then deconvoluted by the Huygens Professional Software.

### Latrunculin A rescue experiment

hTERT-RPE1 control and *RPGR* KO cells were seeded on coverslips (25 mm diameter, thickness #1.5) with a density of 2.1 x 10^4^ cells/cm^2,^ and 6 hours after cell seeding, the DMEM/F12 growth media was supplemented with 0.2 μM Latrunculin A (Santa Cruz, sc-202691) for another 24 hours. The control cells were treated with DMSO. Samples were then immunostained as described above with antibodies against acetylated tubulin, ARL13B, and DAPI. Skin fibroblast cell Latrunculin A treatment was the same as hTERT-RPE1 cells. Airway basal cells were seeded on Transwell supports and differentiated at the air-liquid interface for 2 weeks and switched to ALI medium containing 0.1 μM Latrunculin A for another 2∼3 weeks. After 2 weeks of treatment, samples were collected for cilia beat analysis. After treatment for 3 weeks, cells were harvested to measure ciliation and cilia length.

### Cilia beat analysis

Cilia beat leads to oscillation of signal intensity over time at cilia localized pixels. The contour of the ciliated cells was defined by applying intensity threshold (set as the mean of the whole plot) to the maximum intensity projection plot and a Gauss smoothing. The frequency of oscillations for every pixel of the recording was analyzed by computing the Fourier transform of the corresponding intensity changes over time using the MATLAB fast Fourier transform function (fft()). For each pixel, cilia beat frequency (CBF) was calculated as the frequency where the absolute value of the Fourier transform was maximal. This resulted in a scaled spatial CBF map. To reduce background signal fluctuation noise, we listed all connected pixels in the frequency map using the bwconncomp() function and removed regions with fewer than 144 pixels (*91, 92*). Similarly, we defined regions of interest (ROIs) where the intensity change could reflect the single cell cilia beat frequency manually using the ImageJ plugin ROI Manager. The ROI sizes are variable depending on the actual cell size. Similar FFT process was performed in R to generate the CBFs. For samples with less ciliated cells, we tried to take all cells into account and randomly picked >30 cells in highly ciliated samples. For cilia beat mode analysis, the end-on-view WGA-488 labeled cilia beat videos were analyzed manually. Except the normal beating, four other different types of beat mode can be identified: 1) static cilia; 2) minuscule beat which means cilia beat is restricted with pretty low CBF; 3) tremendous beat which means that cilia beat amplitude is large but there is a loss of power and recovery stroke; 4) uncoordinated beat which means cilia beat in rotationary or/and uncoordinated manner. For each ROI, the cell number for each beat mode was counted, and the ratio was calculated by dividing cell number for each beat mode by the total cell number.

### Single cell sequencing and data analysis

We customized the alignment reference by removing the gene-level annotation for the RPGR gene from the GRCh38 human genome reference obtained from ENSEMBL, and designating each RPGR transcript isoform as an independent gene. A new genome reference was then constructed using Cell Ranger (v7.0.0) (*93*) (the code for this customization is available at https://github.com/Jiayi-Zheng/Build-Gene-Isoform-Reference-for-10X).

Sequence alignment was subsequently performed on a previously published human fetal tracheal scRNA-seq dataset (gestation week 21) (*47*). The dataset was processed and further divided to exclude non-epithelial cells with Seurat (v5.0.0), and VlnPlot() was used to visualize the raw count distribution of RPGR transcripts across various cell types.

### RNA extraction and RT-PCR

The total RNA of cultured human airway MCCs was extracted and reverse-transcribed according to the manufacturer’s protocol (Vazyme, RC112, and R412). The obtained cDNA was then amplified using the forward primer: CAGATGAGGAAGTAGAGATCCCAGAG and reverse primer (*94*): CTCTCCTTCCTCCTTTTCAC. The obtained product was assessed by DNA gel electrophoresis and Sanger sequencing. The amplified band with primers of GAPDH was applied as the loading control.

### TEM

Human nasal cells were scaped and fixed in 2% glutaraldehyde diluted in 100 mM sodium cacodylate, dehydrated with 200 mM D-sucrose dissolved in 100 mM sodium cacodylate. The samples were post-fixed with 1% OsO4, dehydrated with the graded ethanol, and finally embedded in resin. Samples were sliced to a thickness of ∼90 nm and imaged with the FEI Tecnai 20 TEM.

### Statistical analysis

The details of the statistics in this study are specified in the main text, figures, and/or figure legends.

## Supplementary Materials

### Supplementary figures

Fig.S1. Validation of the predicted *RPGR* variants.

Fig.S2. MCCs with RPGR defect presented reduced cilia length and disrupted planar polarity.

Fig.S3. Workflow of single-cell cilia beat frequency analysis.

Fig.S4. Validation of the expression of *RPGR* ORF15 isoform in human airway MCCs.

Fig.S5. RPGR defect doesn’t affect transition zone and axoneme ODAs/IDAs components.

Fig.S6. Apical F-actin meshwork accumulated in the MCCs from RP patients with pathological *RPGR* variants.

Fig.S7. RPGR defect leads to diminished apical Gelsolin.

Fig.S8. Cilia issues for *RPGR* KO hTERT-RPE1 can be rescued with LatA treatment.

Fig.S9. RPGR expression at different developmental stages and the timeline for the LatA rescue experiment.

Fig.S10. LatA treatment partially rescued the affected MCCs with pathological *RPGR* variants.

### Supplementary tables

Table S1. (1) List of patients involved in this study with the disease-associated *RPGR* variants. (2) Disease-associated *RPGR* variants in the patient cohort.

Legend: gDNA: genomic DNA position of variant; ClinVar: database of reported variant with associated phenotype; ACMG: American College of Medical Geneticists variant classification system; GnomAD: control database; Revel, SIFT, MutScore, PhiloP, Splice AI and Pangolin: publicly available software used to determine the pathogenicity of variants.

Table S2. (1) Summary of the airway assessments. Legend and abbreviations: Age is in years, NNO: nasal nitric oxide measurements, CXR: chest X-ray, CT: computed tomography. (2) Summary of the eye phenotypes. Legend and abbreviations: Age in years at the time of test; BCVA: best corrected visual acuity; OD: right eye, OS: left eye, refraction is in diopters; ERG: electroretinogram; RCD: rod-cone dystrophy, when the cases are advanced it is hard to determine; GVF: Goldmann visual field width in degrees (normal width is 120 degrees); HRR: color vision test, result out of 6 plates; CRT: central retinal thickness measured in µm.

Table S3. Summary of all the patients’ cilia characteristics in this study. Table S4. List of antibodies used in this study.

Movie S1. Cilia beat video for MCCs from healthy control, #9, and #28.

Movie S2. Cilia beat video for control and *RPGR* KO MCCs (3 biological replicates)

Movie S3. Defective mucociliary clearance in *RPGR* KO MCCs revealed by the beads propelling experiment

Movie S4. Cilia beat video for control and *RPGR* KO MCCs treated with DMSO and LatA

Movie S5. Cilia beat video for healthy control, #9 MCCs with DMSO treatment, #9 MCCs with LatA treatment, #11 MCCs with DMSO, and #11 MCCs treated with LatA

## Acknowledgment

We thank the patients and their families involved in this study and the healthcare providers. We thank Ting XIE (HKUST) and Shuhuai YAO (HKUST) for sharing cell lines, Randy Yat Choi POON (HKUST) for sharing Caesium 137 gamma radiator, and the Biosciences Central Research Facility (HKUST) for providing the super-resolution microscopy of Zeiss Elyra 7 with Lattice SIM, and the Biosciences Central Research Facility (HKUST-GZ) for providing the STEDYCON STED Super-Resolution Microscope. We also thank students and technicians who contributed to this study’s smaller portions: Alexandra Albulesco, Dan Eddy, Tuo Zhang, Quynh P.H Nguyen, and Alexa Fitzpatrick.

## Author contributions

Y.W., Z.L. designed the experiments and Y.W. performed most experiments. S.D., W.W., and E.H. evaluated the clinical phenotypes and provided the patient samples. E.T., J.M.L. collected the patient samples, performed immunostaining of the fresh nasal cells, and collected the confocal images. B.L. developed methods and analyzed the cilia beat data. J.C. generated the RPGR KO hTERT-RPE1 cells. V.M. and H.F. participated in the early studies of the project. Y.W. and Z.L. wrote the manuscript. Z.L., S.D., E. H, and E.T., revised the manuscript.

## Data and materials availability

All data related to this project are present in the paper or the Supplementary Materials.

