## supplementary figures for "RPGR regulates motile cilia by interfering with actin dynamics"

### Supplementary Materials

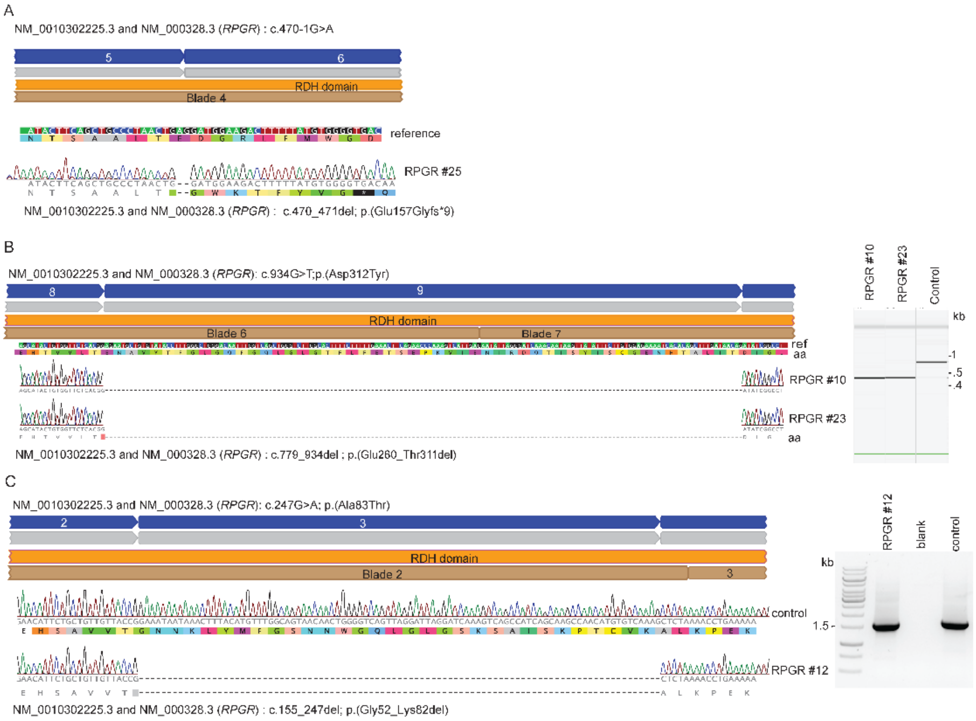


**Fig. S1. Validation of the predicted *RPGR* variants.** RT-PCR and Sanger sequencing of PCR product from control and patient-derived cell lines.

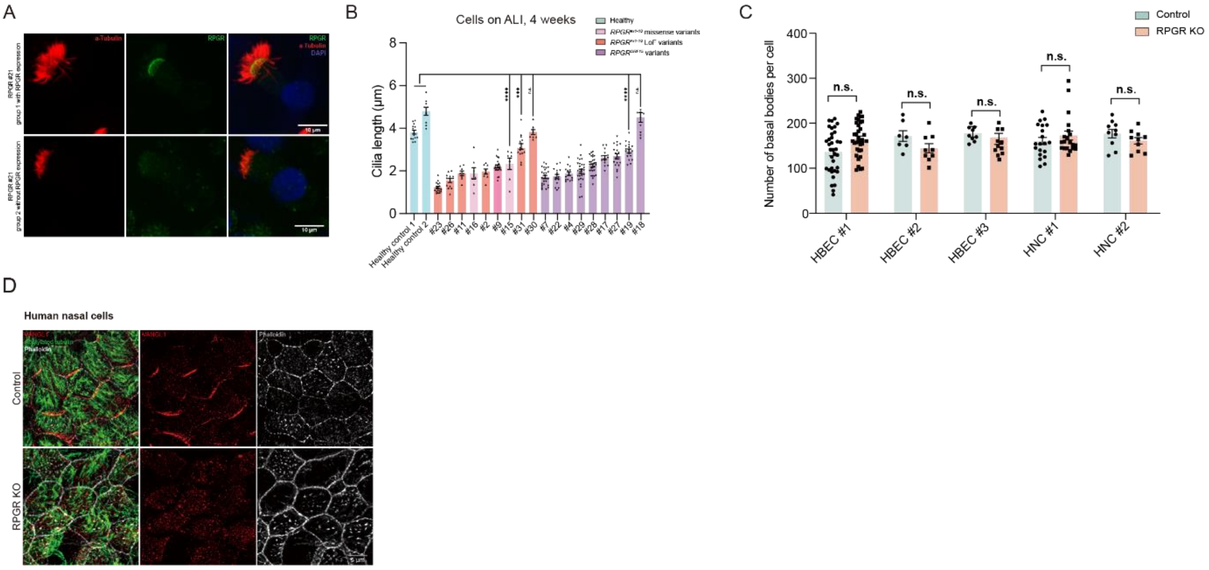


**Fig. S2. MCCs with RPGR defect presented reduced cilia length, and disrupted planar polarity. (A)** Immunostaining of RPGR and cilia marker a-tubulin showed that MCCs for RPGR #21 with RPGR expression presented normal cilia length, and those without RPGR expression presented decreased cilia length. Scale bar 10 µm. **(B)** Cilia length measurements for the MCCs from both controls and RP patients cultured at the air-liquid interface for 4 weeks. n>6 cells per sample. **(C)** The number of basal bodies was unaffected for all 4 RPGR KO biological replicates. **(D)** Immunostaining of planar polarity Vangl1, cilia marker acetylated tubulin, and Phalloidin showed Vangl1 mislocalization for *RPGR* KO HNCs. Scale bar 5 µm. All data are presented as average ± SEM. n.s. no significance, *, $p < 0.05$, **, $p < 0.01$, ***, $p < 0.001$, ****, $p < 0.0001$ by two-tailed t-test (B), or two-way repeated ANOVA followed by Sidak's post hoc test (C).

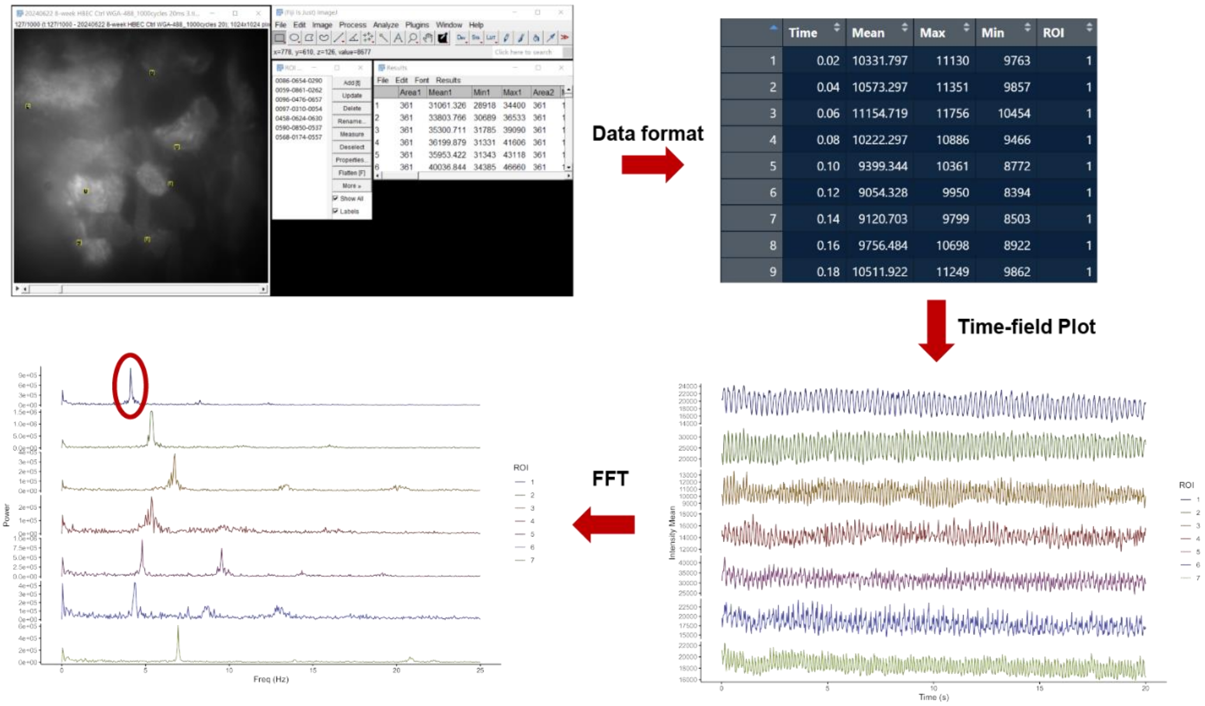


**Fig. S3. Workflow of single-cell cilia beat frequency analysis.** The peak frequency generated from the manually acquired intensity spectrum by fast Fourier transform is designated as the cilia beat frequency of the cell.

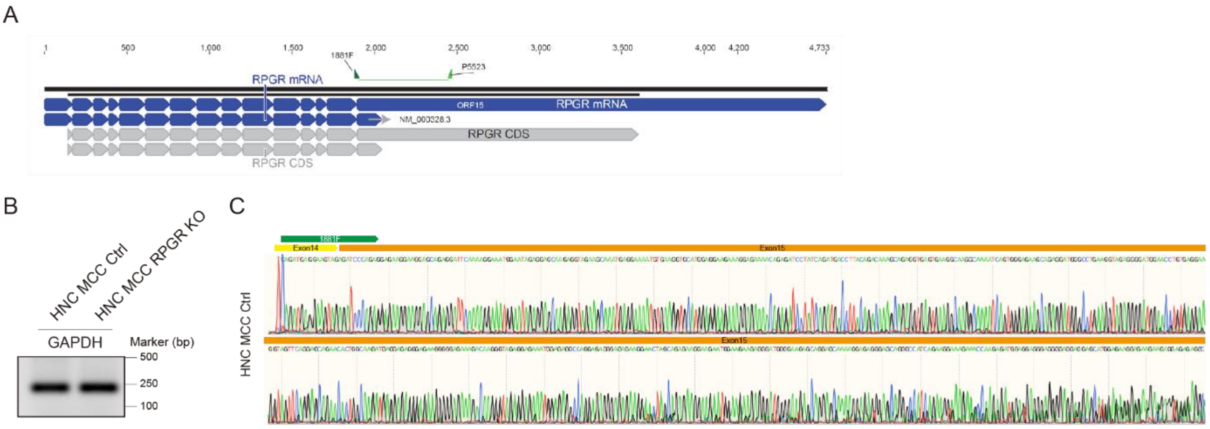


**Fig. S4. Validation of the expression of *RPGR* ORF15 isoform in human airway MCCs. (A)** RT-PCR primer design for RPGR ORF15 isoform. **(B)** Loading control for RT-PCR shown in Fig. 2J. **(C)** The amplified RT-PCR band (Fig. 2J) was purified and validated by Sanger sequencing.

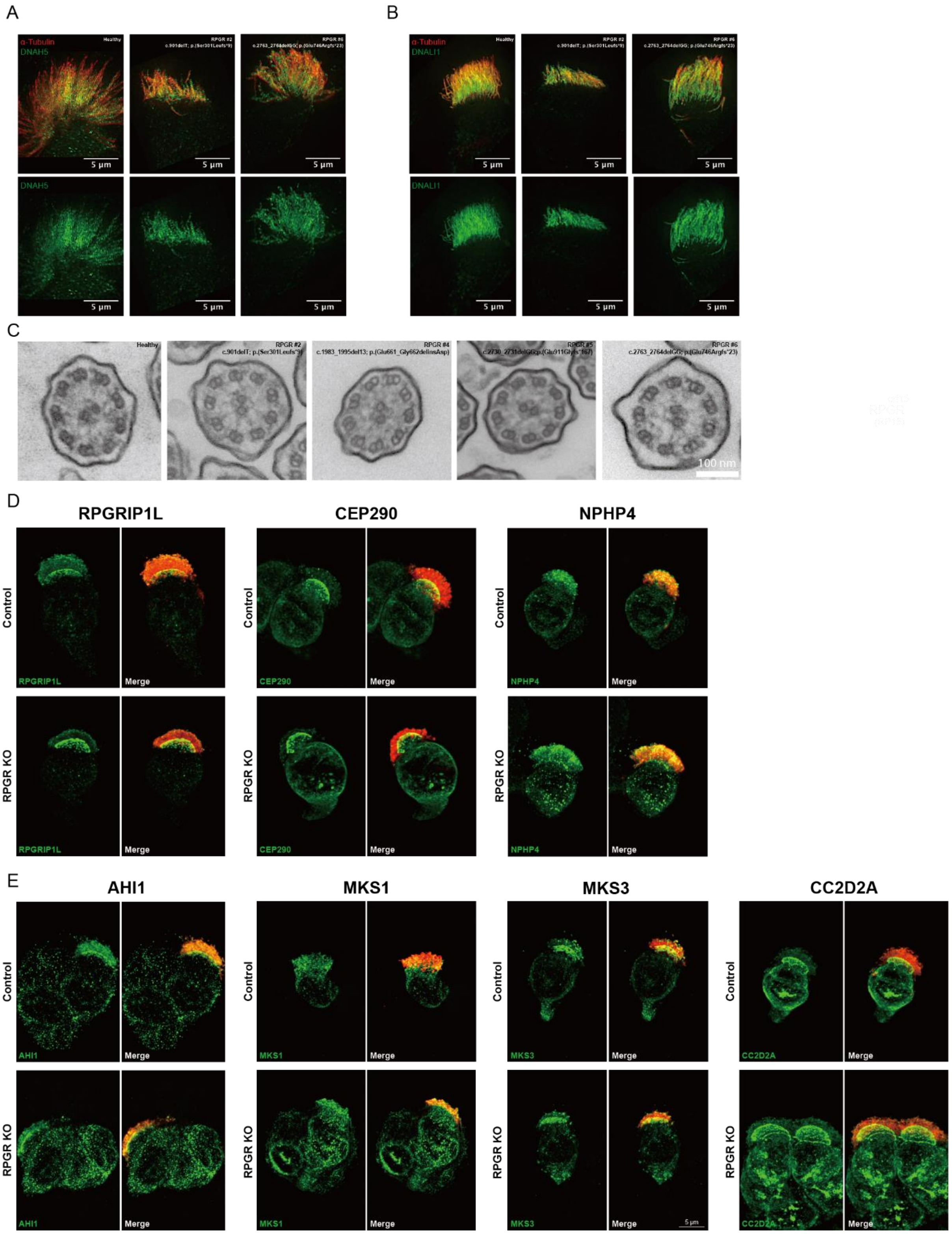


**Fig. S5. RPGR defect doesn't affect transition zone and axoneme ODAs/IDAs components. (A)** Immunostaining of ODA marker DNAH5 and cilia marker a-tubulin showed that the MCCs with *RPGR* and *RPGR*$^{ORF15}$ variants had intact ODA components. Scale bar 5 μm. **(B)** Immunostaining of IDA marker DNLI1 and cilia marker a-tubulin showed that the MCCs with *RPGR and RPGR*$^{ORF15}$ variants had intact IDA components. Scale bar 5 μm. **(C)** TEM showed the normal ciliary ultrastructure for human MCCs with different *RPGR* variants.

Scale bar 100 nm. **(D)** Immunostaining of RPGR interactors RPGRIP1L, CEP290, or NPHP4 and cilia marker acetylated tubulin showed that MCCs with RPGR defect didn't affect their TZ distributions. Scale bar 5 μm. **(E)** Immunostaining of cilia maker acetylated tubulin and transition zone components AHI1, MKS1, MKS3, or CC2D2A showed that RPGR defect preserved the normal transition zone structure. Scale bar 5 μm.

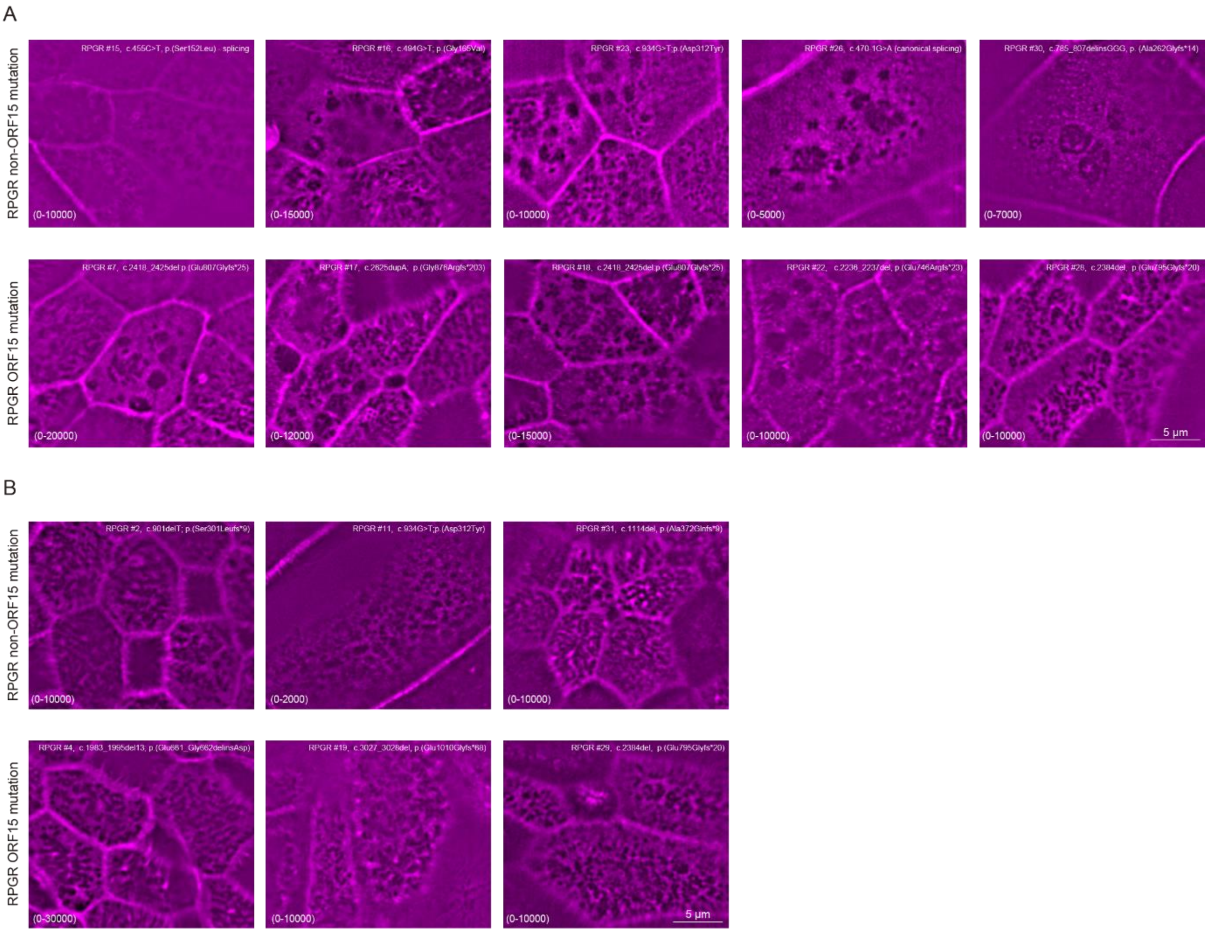


**Fig. S6. Apical F-actin meshwork accumulated in the MCCs from RP patients with pathological *RPGR* variants. (A)** The apical surface was found to be covered with F-actin meshwork for the mature MCCs from RP patients with *RPGR* pathological variants. Scale bar 5 μm. **(B)** The mature MCCs of some RP patients (6/19) didn't show obvious F-actin meshwork. Scale bar 5 μm.

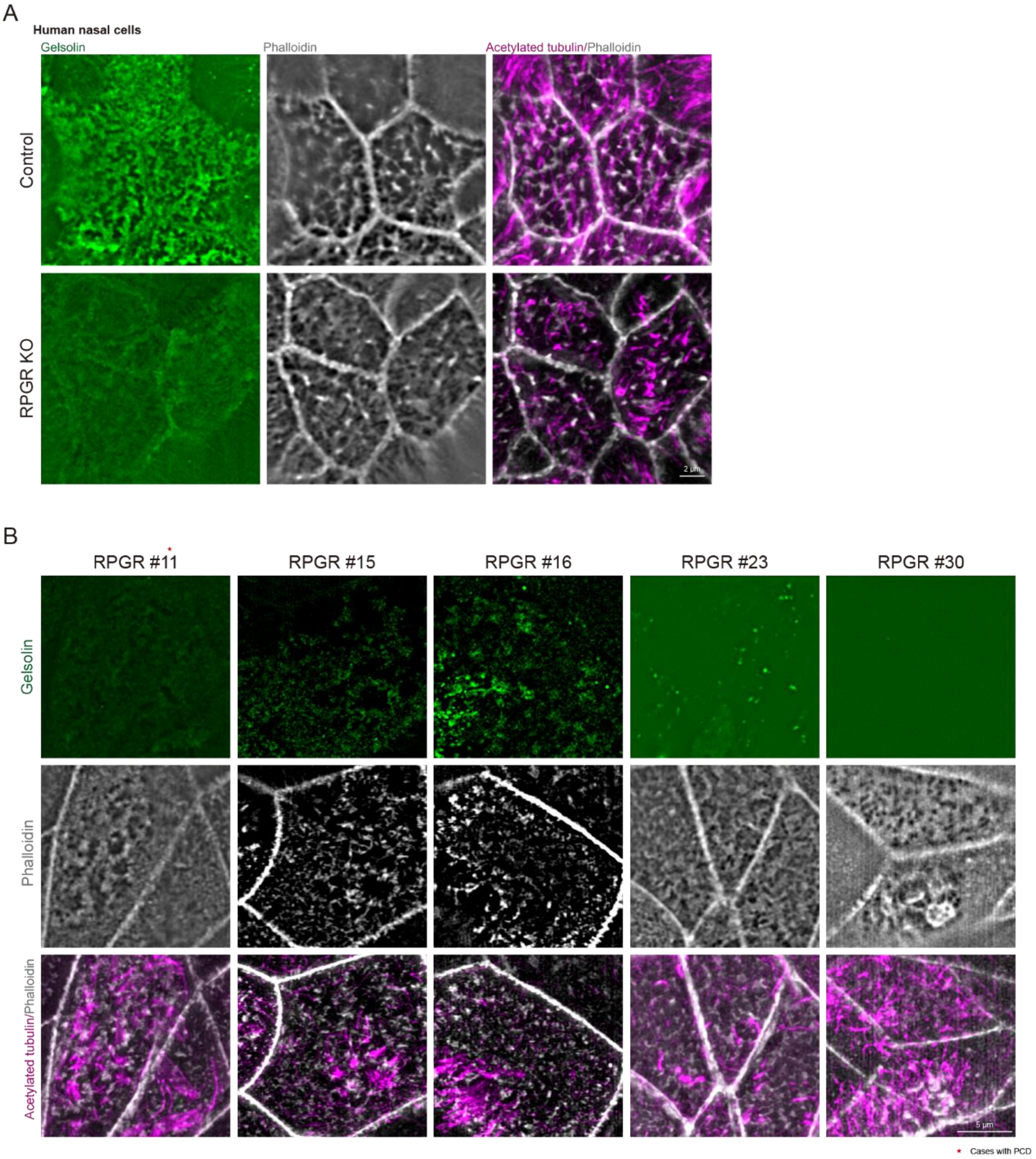


**Fig. S7. RPGR defect led to diminished apical Gelsolin. (A)** 4-week *RPGR* KO HNC MCCs showed diminished apical gelsolin. Scale bar 2 µm. **(B)** The diminished apical gelsolin was observed in the 4-week MCCs from RP patients with *RPGR* pathological variants. Scale bar 5 µm.

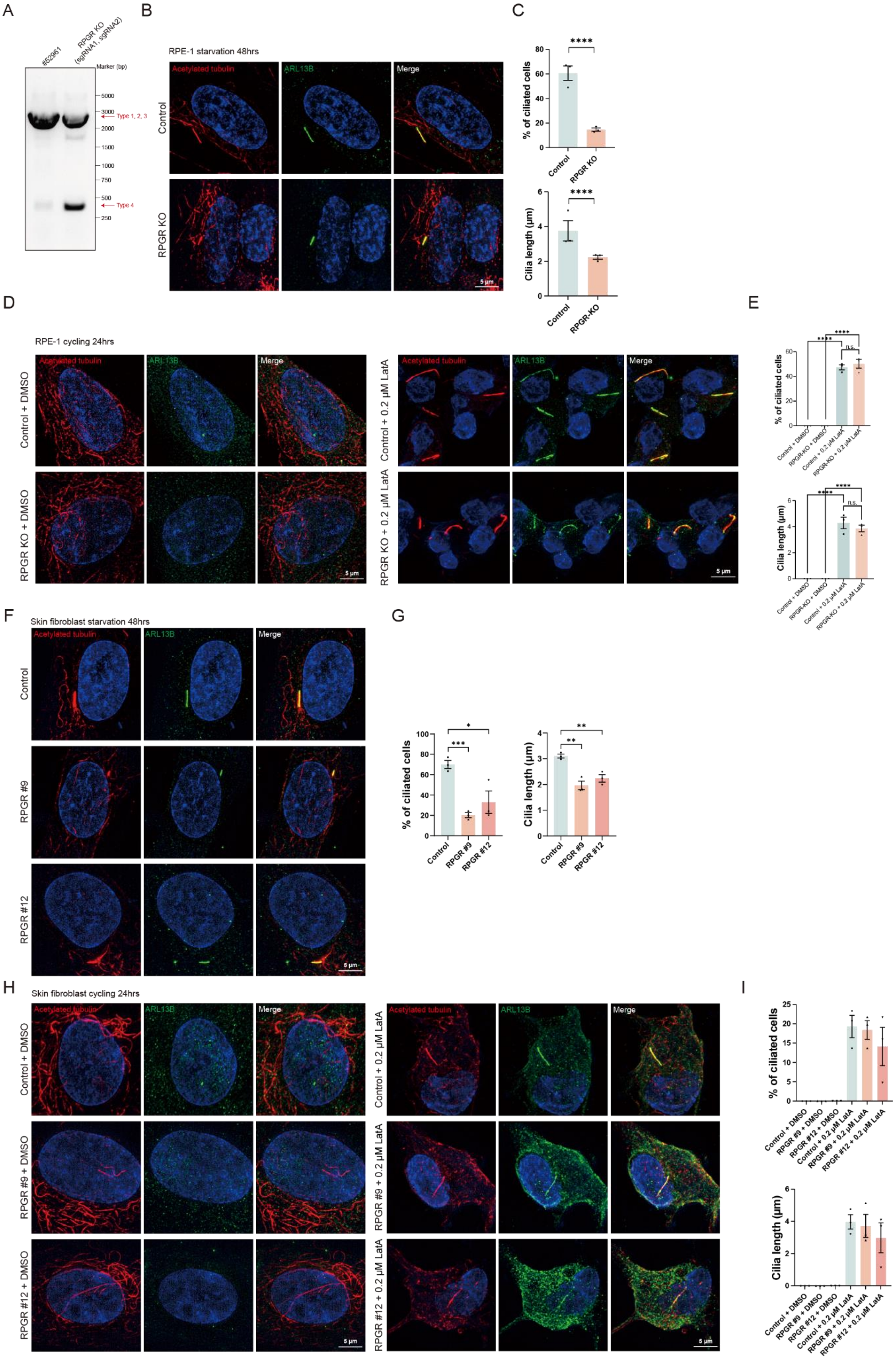
A
B
C
#52961
RPGR KO (sgRNA1, sgRNA2)
Marker (bp)
5000
3000
2000
1500
1000
750
500
250
Type 1, 2, 3
Type 4
RPE-1 starvation 48hrs
Acetylated tubulin
ARL13B
Merge
Control
RPGR KO
5 µm
% of ciliated cells
Cilia length (µm)
RPGR-KO
D
E
RPE-1 cycling 24hrs
Control + DMSO
RPGR KO + DMSO
Control + 0.2 µM LatA
RPGR KO + 0.2 µM LatA
n.s.
F
G
Skin fibroblast starvation 48hrs
RPGR #9
RPGR #12
H
I
Skin fibroblast cycling 24hrs
RPGR #9 + DMSO
RPGR #12 + DMSO
RPGR #9 + 0.2 µM LatA
RPGR #12 + 0.2 µM LatA

**Fig. S8. Cilia issues for *RPGR* KO hTERT-RPE1 can be rescued with LatA treatment. (A)** DNA gel showed the high efficiency of *RPGR* KO in hTERT-RPE1. **(B, C)** Loss of RPGR in hTERT-RPE1 cells led to reduced ciliation and short cilia length. Scale bar 5 μm. **(D, E)** The ciliation and cilia length issues of *RPGR* KO in hTERT-RPE1 cells were rescued by Lat A treatment. Scale bar 5 μm. **(F, G)** Skin fibroblasts with *RPGR* pathological variants showed reduced ciliation and cilia length. Scale bar 5 μm. **(H, I)** Cilia issues of patient fibroblast cells caused by *RPGR* variants can be rescued by LatA. Scale bar 5 μm. All data are presented as average ± SEM. n.s. no significance, *, $p < 0.05$, **, $p < 0.01$, ***, $p < 0.001$, ****, $p < 0.0001$ by two-tailed t-test (C, F).

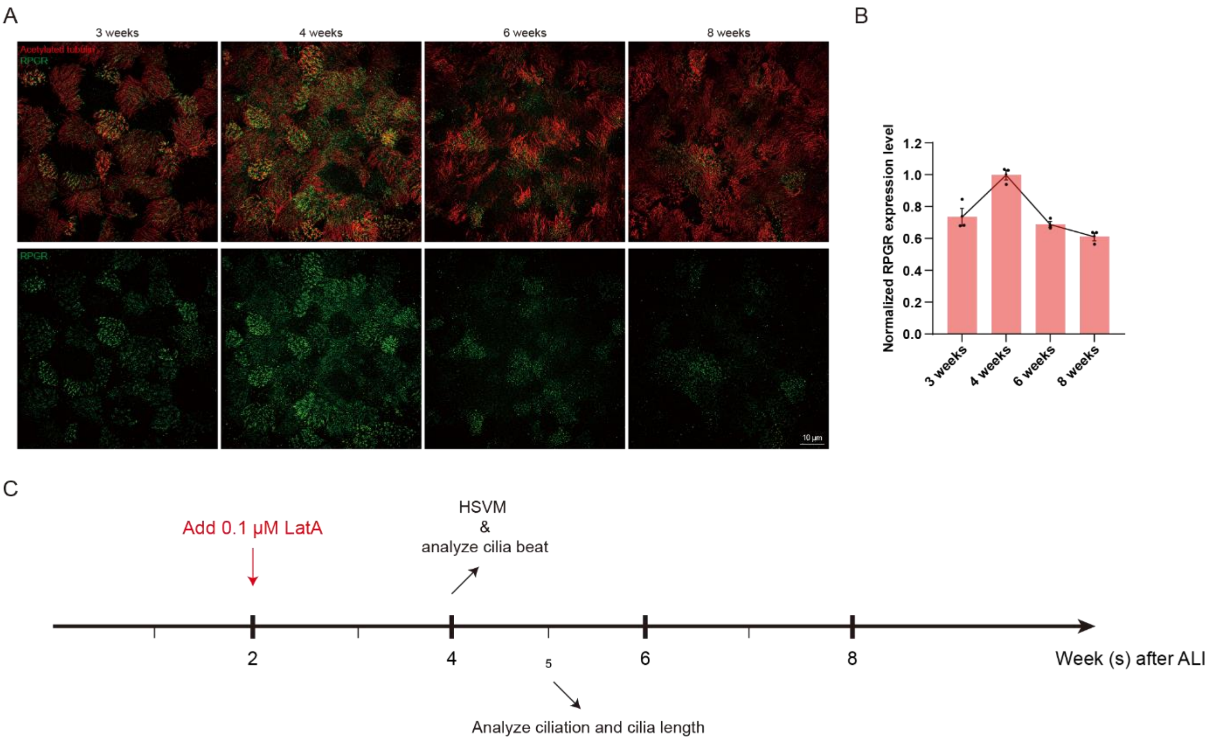


**Fig. S9. RPGR expression at different developmental stages and the timeline for LatA rescue experiment. (A)** Immunostaining of RPGR and cilia marker acetylated tubulin at different developmental stages. Scale bar 10 μm. **(B)** Expression of RPGR peaks at ALI 4 weeks. **(C)** Lat A was added at ALI 2 weeks, cilia beat was observed at 4 weeks, and ciliation and cilia length were analyzed at 5 weeks. All data are presented as average ± SEM.

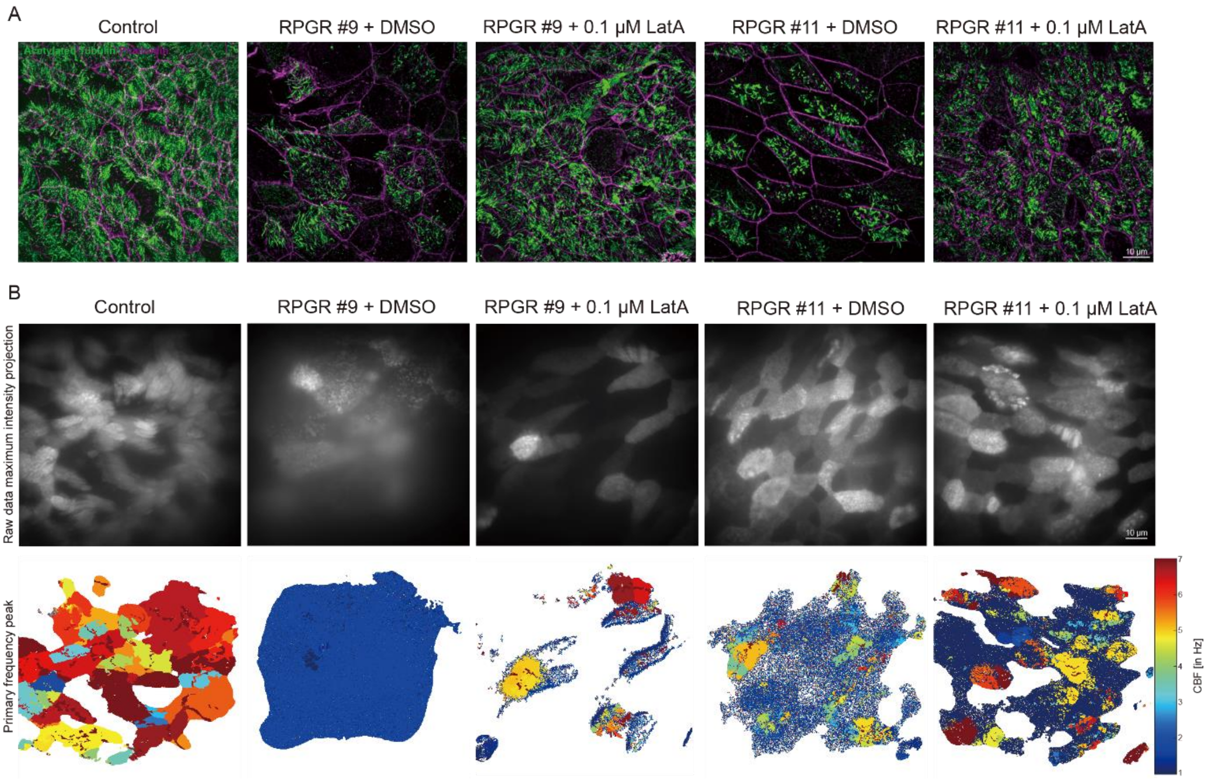


**Fig. S10. LatA treatment partially rescued the affected MCCs with pathological variants.** **(A)** Immunostaining with anti-acetylated tubulin antibodies and phalloidin-Alexa647 showed that LatA can increase the ciliation and cilia length. Scale bar 10 μm. **(B)** Motile cilia beat analysis showed LatA treatment can partially rescue CBF. Scale bar 10 μm.
